# Analysis of a shark reveals ancient, Wnt dependent, habenular asymmetries in jawed vertebrates

**DOI:** 10.1101/2023.10.17.562666

**Authors:** Maxence Lanoizelet, Léo Michel, Ronan Lagadec, Hélène Mayeur, Lucile Guichard, Valentin Logeux, Dany Séverac, Kyle Martin, Christophe Klopp, Sylvain Marcellini, Hector Castillo, Nicolas Pollet, Eva Candal, Mélanie Debiais-Thibaud, Catherine Boisvert, Bernard Billoud, Michael Schubert, Patrick Blader, Sylvie Mazan

## Abstract

The origin of left-right asymmetries in the vertebrate habenula remains largely unknown. Using a transcriptomic approach, we show that in a cartilaginous fish, the catshark *Scyliorhinus canicula*, habenulae exhibit marked asymmetries both in their medial and their lateral component. Comparisons across gnathostomes suggest that asymmetries in the catshark lateral habenulae reflect an ancestral gnathostome trait, independently lost in tetrapods and neopterygians. Analysis of the mechanisms underlying their formation highlights an essential role of Wnt signaling. Wnt activity is submitted to a dynamic, asymmetric regulation during habenula development, with a Nodal dependent left repression at a stage when precursors for lateral habenulae have exited cell cycles. Pharmacological treatments during this time window reveal that Wnt signaling promotes lateral right neuronal identities in the right lateral habenula, while its repression by Nodal in the left one promotes lateral left neuronal identities. Based on comparisons with the zebrafish and the mouse, we propose that habenular asymmetry formation and diversification in gnathostomes involve the same developmental logic, relying on a conserved temporal regulation of neurogenesis, shaping neuronal identities on both sides, and its modification by a dynamic Wnt activity, right-restricted in the ancestral state and prone to variations in time and space during evolution.

## INTRODUCTION

Habenulae are bilateral epithalamic structures, present in all vertebrates. They appear as a key node in conserved neuronal circuits connecting various forebrain areas to midbrain nuclei, and integrate information from multiple sources, including sensory organs and corticolimbic areas, to regulate complex behavioral, cognitive and emotional responses^1–4^. A basic architecture of this structure into two domains (medial and lateral in mammals, dorsal and ventral in teleosts), exhibiting distinct molecular properties and connectivity patterns, is known to be conserved across gnathostomes^5,6^. Molecular characterizations, including detailed single cell-RNA-sequencing maps highlight similarities between the mouse medial and the zebrafish dorsal habenulae, as well as between the mouse lateral and the zebrafish ventral habenulae^7,8^, but more precise comparisons of neuronal populations across vertebrates remain difficult. A unique feature of the habenulae in the vertebrate brain is that it displays asymmetries between the left and the right sides in many species including human^9,10^. Their biological significance has started to emerge in the zebrafish, the reference model for analyses of both their roles and mechanisms of formation^11^. In this species, sensory cues are differentially processed between the right and left habenulae, and asymmetries have been shown to impact important adaptive responses, such as exploratory and food-seeking behaviors, light preference or responses to fear^12–15.^. Despite the evolutionary relevance of these functions depending on ecological context, the origin and mode of diversification of habenular asymmetries across vertebrates remain unclear. Morphological characterizations suggest that they considerably vary in nature and degree across vertebrates (size, cellular or nuclei organisation, connectivity pattern)^10^. In line with this complexity, more detailed molecular characterizations, essentially focused on teleosts thus far, highlight complex presence/absence patterns of asymmetric traits, suggestive of a rapid drift in this taxon^16,17^. However, whether habenular asymmetries in teleosts reflect the vertebrate or gnathostome ancestral state is unclear, since they may employ highly divergent mechanisms for their formation^18^. In the zebrafish, habenular asymmetries are restricted to the dorsal habenula (dHb), and they consist of different relative proportions between two subdomains, respectively occupying lateral (dHbl) and medial locations (dHbm) and prevailing on the left and on the right sides^19–21^. Analyses of the mechanisms controlling their formation have highlighted a pivotal role for the Wnt signaling pathway. Its activity is submitted to a finely tuned temporal control during habenular development and it controls distinct successive processes, including the specification of the ventral habenula, the regulation of the temporal control of neurogenesis and the choices of neuronal identities in the differentiating dorsal habenula^22–25^. Its repression on the left side via an interaction with the adjacent parapineal during a discrete time window results in the elaboration of habenular asymmetries, and the promotion of right-sided neuronal identities in the right dorsal habenula^24^. Analyses of two non-conventional models with marked habenular asymmetries, the lamprey and the catshark, suggest that this mechanism does not reflect the ancestral vertebrate state, which was likely independent from the parapineal and instead involved an early left-restricted diencephalic activity of Nodal, dispensable in the zebrafish^18^. To clarify the mode of evolution of habenular asymmetries across gnathostomes (jawed vertebrates), we have focused on the catshark *Scyliorhinus canicula*, a member of chondrichthyans (cartilaginous fishes), which are the sister group of osteichthyans (bony fishes and tetrapods) and therefore provide a relevant reference to explore the origin of habenular asymmetries^26^. We find that habenulae in this species display a highly asymmetric organization, retaining ancient asymmetries of jawed vertebrates, that have been lost in the mouse and zebrafish. We also propose a conserved regulatory logic for asymmetry formation, involving interactions between a conserved temporal control of neurogenesis and a more flexible temporal and spatial regulation of Wnt signaling, which could account for the evolvability of habenular asymmetries across vertebrates.

## RESULTS

### Identification of differentially expressed genes between developing left and right catshark habenulae

To obtain an unbiased characterization of molecular asymmetries in catshark habenulae, we conducted a transcriptomic comparison between developing left and right habenulae. We focused on stage 31, when a large fraction of the organ is differentiated but asymmetric pools of neural progenitors persist^27^. Statistical analysis of 362 million read pairs obtained from three replicates of left versus right habenulae explants led to the identification of 614 differentially expressed gene models (373 left-enriched and 241 right-enriched), of which 538 were annotated as protein coding (Fig. 1a; Supplementary Table 1). Functional annotation of left- and right-enriched gene lists shows that they share an over-representation of GO (Gene Ontology) terms related to neurogenesis, neuronal differentiation, the formation of neuronal projections and trans-synaptic signaling, as expected for a differentiating brain structure (Supplementary Fig.1a,b; Supplementary Table 1). GO terms selectively over-represented in either one of these two lists were also detected. They include terms related to organismal responses known to be regulated by habenulae, such as the perception of pain and behavioral fear responses, and to signaling pathways known to be active in habenulae (Wnt, opioid receptor, and ionotropic glutamate receptor signaling) (Supplementary Fig.1c-d). Altogether, these data suggest differential functional specializations of the catshark left and right habenulae.

### Catshark habenulae exhibit an asymmetric, lateral to medial subdomain organization

To validate asymmetries identified by the transcriptomic analysis and to characterize their spatial organization, we performed chromogenic *in situ* hybridization (ISH) on sections of stage 31 habenulae for a total of 39 genes, selected from the lists of asymmetrically expressed genes (Fig. 1a). Regionalized signals were observed for most of these genes, with three of them showing restricted expressions in territories consisting of neural progenitors (Supplementary Fig.2h)^27^, and twenty-nine in differentiating subdomains of the habenulae, all consistent with the expression lateralities predicted by the transcriptomic analysis (Supplementary Fig.2b-g). The twenty-nine genes can be classified into five broad categories, each characterized by largely overlapping expression profiles restricted to (i) a left lateral territory (Supplementary Fig.2b), (ii) a right lateral one (Supplementary Fig.2f), (iii) a broad bilateral medial one larger on the left than on the right (Supplementary Fig.2d), (iv) a left-restricted subdomain of this medial territory (Supplementary Fig.2c) and a right-enriched subdomain of the medial territory (Supplementary Fig.2e). More detailed analyses focusing on markers of each of these territories (*ScSox1*, *ScPde1a*, *ScKctd12b*, *ScEnpp2*, and *ScProx1*; Fig.1; Supplementary Fig.3 and 4) highlight an organization of stage 31 catshark habenulae into three complementary broad subdomains, two lateral ones, referred to as Left- and Right-LHb hereafter, expressing *ScSox1* on the left and *ScProx1* on the right, and a broad bilateral *ScKctd12b*-positive medial one, referred to as MHb, including a previously described left-restricted anterior component^18^ (Fig. 1b2,d,f-k,n,o,q,s,u; Supplementary Fig.3b,d,f; Supplementary Fig.4b,d,f). *ScSox1* and *ScProx1* expressions are confined, respectively, to the left and the right lateral habenulae, except for a small contralateral neuronal population at posterior-most levels of the habenulae (Fig.i,k; Supplementary Fig.3b5,f4 and Supplementary Fig.4b4,f3). The bilateral medial territory (MHb) is itself partitioned into an internal component and an external one, adjacent to Left-LHb and Right-LHb (Fig. 1c,e,r,t). The external MHb component is organized into two discrete abutting territories, an anterior, left restricted *ScPde1*-positive territory (Fig. 1c,l,r; Supplementary Fig.3c,4c) and a *ScEnpp2*-positive one, occupying the whole external part of MHb on the right and its posterior part on the left (Fig. 1e,m,t; Supplementary Fig.3e,4e). To assess whether these asymmetries reflect transient ones in the differentiating habenulae or definitive neuronal identities of the functional organ, we investigated their maintenance in feeding juveniles collected after exhaustion of their yolk reserves (Supplementary Fig.5). The broad expression characteristics and asymmetries of *ScSox1*, *ScPde1a*, *ScKctd12b*, *ScEnpp2*, and *ScProx1* are very similar to those observed at stage 31 (Supplementary Fig.5b-j), except for an expansion of the *ScEnpp2* signal to the internal component of MHb (Supplementary Fig.5e). Altogether, these results highlight a highly asymmetric organization of the catshark habenulae, both in its lateral and medial subdomains.

**Figure 1.**
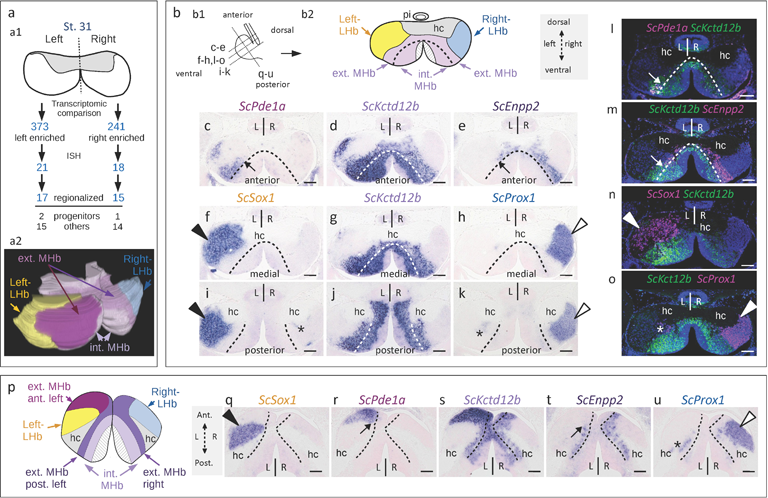
Developing catshark habenulae harbor major asymmetries both in lateral and medial habenulae. **a** Schemes showing the experimental strategy for the characterization of molecular asymmetries in catshark habenulae (a1) and the resulting 3D organization at stage 31 (a2; anterior to the left, dorsal to the top). **b** Schemes showing a left lateral view of catshark stage 31 habenulae, with section planes and levels indicated by dotted lines (b1), and the subdomain organization observed on a transverse section at a medial level (b2). **c-o** Transverse sections (dorsal to the top of each panel) after *in situ* hybridization (ISH) with probes for *ScSox1* (f,i), *ScPde1a* (c), *ScKctd12b* (d,g,j), *ScEnpp2* (e), *ScProx1* (h,k), and after fluorescent double ISH with probes for *ScPde1a*/*ScKctd12b* (l), *ScKctd12b*/*ScEnpp2* (m), *ScSox1*/*ScKctd12b* (n), and *ScKctd12b*/*ScProx1* (o). For fluorescent double ISH, signals for *ScKctd12b* are shown in green and those for *ScPde1a*, *ScEnpp2*, *ScSox1*, and *ScProx1* in magenta. **p** Scheme showing the subdomain organization of stage 31 habenulae, observed on a horizontal section at a medial level. **q-u** Horizontal sections (anterior to the top of each panel) after ISH with probes for *ScSox1* (q), *ScPde1a* (r), *ScKctd12b* (s), *ScEnpp2* (t), and *ScProx1* (u). Sections (c-e), (f-h), and (i-k) were obtained from the same embryo, same for (q-u). Black and white arrowheads respectively point to major lateral territories of *ScSox1* and *ScProx1*, and asterisks show contra-lateral minor posterior territories. Dotted lines in (c-m,q-u) delimit external and internal subdomains of the medial habenula, as inferred from the inner boundaries of *ScPde1a* and *ScEnpp2* territories. Thin arrows in (c,e,l,m,r,t) point to the boundary between the complementary territories of *ScPde1a* and *ScEnpp2* within the external MHb subdomain. Color code in (b2): yellow, Left-LHb; light purple, MHb; blue, Right-LHb; hatched, pseudo-stratified neuroepithelium containing neural progenitors. Color code in (a2,p): yellow, Left-LHb; light purple, internal MHb; magenta, anterior-left component of external MHb; dark purple, right plus posterior-left components of external MHb; hatched, neural progenitors. Abbreviations: ant., anterior; post., posterior; ext., external; int., internal; MHb, medial habenula; LHb, lateral habenula; hc, habenular commissure; pi, pineal stalk; L, left; R, right. Scale bar=100μm.

### Comparisons between the catshark, the mouse and the zebrafish highlight similarities in subdomain signatures, without asymmetry conservations

To identify conserved features of habenular asymmetry or subdomain organization across gnathostomes, we initially focused on habenular expression of mouse and zebrafish orthologs of markers of the five major habenular territories identified in the catshark. Orthologs of *Kctd8/12a/12b* were included in this analysis (Supplementary Table 2; Supplementary Fig.6). In the mouse, this analysis was based on profiles published in the habenulae, including systematic searches in the Allen Brain Atlas^28^. Most mouse orthologs of catshark MHb markers are expressed in the mouse medial habenula, including *Spon1* and *Trhde*, restricted to the anterior left external component of MHb in the catshark and to a lateral subdomain of the medial habenula in the mouse (Supplementary Table 2; Supplementary Fig.6a-c,k). Two of the four gene signatures of Left-LHb in the catshark (*Pcdh17* and *Ntng2*) share a bilateral expression in mouse lateral habenulae (Supplementary Fig.6d-e), an expression site also observed for a *Sox1* reporter in *Sox1*-*LacZ* knock-in mice^29^. Mouse orthologs of catshark Right-LHb markers show various expression profiles spanning the ventral medial habenulae, lateral habenulae and adjacent thalamic nuclei (Supplementary Fig.6f-j), with three of them sharing a highly specific expression territory in the paraventricular nucleus of the thalamus, ventral to the lateral habenulae (*Rerg*, *Prox1*, and *Stxbp6*: Supplementary Fig.6h-j). For comparisons with the zebrafish, we examined the presence of orthologs of catshark territory markers in gene signatures of single-cell RNA-seq gene clusters mapped to larval or adult habenulae^8^ (Supplementary Table 2). Orthologs of three asymmetrically expressed catshark MHb markers (*Spon1*, *Kctd8*, and *Kctd12a*) also display asymmetric expressions in zebrafish dHb, albeit without consistent conservation of asymmetry laterality. Conversely, with one exception (*prox1a*, part of a dorsal and right-enriched cluster), zebrafish orthologs of catshark LHb markers (*Sox1*, *Ak5*, *Kiss1*, *Rerg*, *Prkcq*, and *Gng14*) are identified as gene signatures of two bilaterally symmetric ventral cell clusters of zebrafish larval habenula cell clusters, Hb11 for zebrafish *sox1a/b*, Hb15 (or related adult clusters) for the other five genes. Taken together, these data support the conservation in the catshark of the subdivision of habenulae in two components (dorsal/medial and ventral/lateral). Intriguingly, they highlight similarities of catshark left-LHB with the mouse lateral habenulae, and of catshark right-LHb with a major zebrafish ventral cell cluster.

### Conserved asymmetry patterns can be detected between chondrichthyans, a dipneust, and a polypterid in the lateral/ventral habenula

The similarities of catshark Left-and Right-LHB respectively with the mouse lateral and the zebrafish ventral habenulae may reflect convergences or taxon-specific diversifications of lateral/ventral habenulae, unrelated to ancestral traits of habenula organization. To test their evolutionary relevance, we used a phylogenetic approach, aimed at reconstructing habenular organization at major gnathostome nodes, taking the catshark as reference. For this analysis, we selected five species for their phylogenetic position in major gnathostome phyla: an holocephalan, the elephant shark *Callorhinchus milii*, two non-teleostean actinopterygians, the reedfish *Erpetoichthys calabaricus* (bichir) and the spotted gar *Lepisosteus oculatus*, and two sarcopterygians, the lungfish *Protopterus annectens* and the frog *Xenopus tropicalis*. In each one of these species, we carried out ISH analyses on transverse sections of habenulae from juveniles/tadpoles (frog), for orthologs of markers of catshark Left-LHb (*Sox1, Ntng2*, and *Pcdh17*), MHb (*Kctd8, Kctd12a*, and *Kctd12b*) and Right-LHb (*Prox1* and *Kiss1*) (Fig.2; Supplementary Fig.7-11). Specific habenular expression was observed in all cases, except for *Kctd12a* in the lungfish, and *Kiss1* in the lungfish and the frog, whose expression remained very weak or undetectable by ISH. In all species analyzed, profiles of members of the *Kctd8*/*12a*/*12b* family vary between paralogs and species, but at least one paralogue always defines a medial or dorsal habenular territory (Fig. 2g-l; Supplementary Fig.7e-g; Supplementary Fig. 8f,i-j; Supplementary Fig.9d-f; Supplementary Fig.10e,f; Supplementary Fig.11f,h-j). With a few exceptions (such as the expansion of elephant shark *Kiss1* and *Pcdh17*, reedfish *Prox1*, and spotted gar *Ntng2* to various MHb/dHb subterritories), orthologs of catshark Left- and Right-LHb markers are expressed within lateral or ventral territories (Fig. 2b-f,n-r; Supplementary Fig.7b-d,h-i; Supplementary Fig. 8b-d,g-h; Supplementary Fig.9g-k; Supplementary Fig.10b-d,g; Supplementary Fig.11b-e,g). However, the labeled territories and asymmetry patterns markedly differ between species. As in the catshark, the elephant shark and the reedfish show left- and right-restricted lateral/ventral territories, respectively co-expressing *Sox1/Pcdh17/Ntng2* and *Prox1/Kiss1* (Fig. 2b,c,n,o; Supplementary Fig.7b-d,h-i; Supplementary Fig. 8b-e,g-h). Similarly, in the lungfish, *Sox1*, *Pcdh17*, and *Ntng2* share a strong left-restricted signal, adjacent to *Kctd8/12b* positive territories. Furthermore, a more discrete lateral *Prox1*-positive cell population lining the habenular commissure is observable on the right side, in a zone devoid of *Kctd8/12* expression (Fig. 2e,q; Supplementary Fig.10b-d,g). In contrast, expression of orthologs of both catshark Left-and Right-LHb markers are bilateral in the spotted gar and the frog, albeit with slightly different relative locations between the two species. In the spotted gar, *Pcdh17/Sox1/Ntng2* share an anterior expression territory, complementary to the *Prox1/Kiss1*-positive one within the left and right ventral habenulae (Fig. 2d,p; Supplementary Fig.9b-c,g-k). In the frog, their expression is also observed both in the left and right habenulae, adjacent and lateral to the domain of *Kctd8/12b* (Fig. 2f; Supplementary Fig11b-e), with *Prox1* being expressed at more ventral and posterior levels (Fig. 2r; Supplementary Fig.11g).

**Figure 2.**
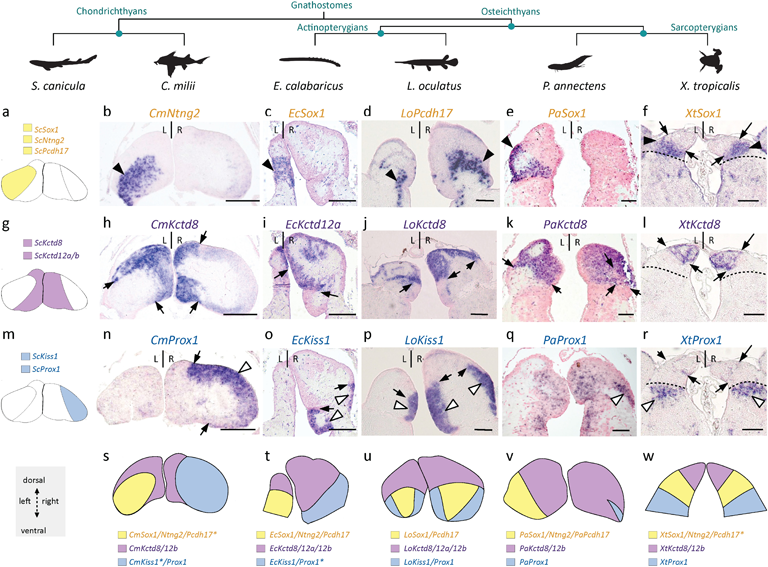
Asymmetries related to those observed in catshark lateral habenulae are detected in dipneusts and polypterids, but not neopterygians and tetrapods. **a,g,m** Schemes showing the asymmetric organization of the catshark habenulae, with the reference signature markers of Left-LHb (a, yellow), MHb (g, purple) and Right-LHb (m, blue) considered for cross-species comparisons. **b-f,h-l,n-r** Transverse sections of habenulae in elephant shark (*Callorhinchus milii*: b,h,n), reedfish (*Erpetoichthys calabaricus*: c,i,o), spotted gar (*Lepisosteus oculatus*: d,j,p), lungfish (*Protopterus annectens*: e,k,q) and western clawed frog (*Xenopus tropicalis*: f,l,r) juveniles, following ISH with probes for orthologs of catshark markers for Left-LHb (b-f), MHb (h-l), and Right-LHb (n-r). Dorsal is to the top in all panels and probe identity is indicated on each view. Black and white arrowheads in (b-f) and (n-r) point to territories respectively restricted to the left and to the right in the elephant shark, the reedfish and the lungfish, similar to the catshark, but bilateral in the spotted gar and the frog. Black arrows indicate the boundary between dorsal/medial and ventral/lateral habenulae. Dotted lines in (f,l,r) delineate the boundary between the lateral *Sox1* habenular territory and adjacent *Prox1* territory in the frog. **s-w** Schemes showing territories related to catshark Left-LHb (yellow), MHb (purple), and Right-LHb (blue) in the elephant shark (s), the reedfish (t), the spotted gar (u), the lungfish (v) and the frog (w), based on the set of signature markers analyzed. Only markers supporting these relationships are indicated for each species, with those also expressed in additional territories labeled by an asterisk (see expression details in Supplementary Fig.7-11). Abbreviations: same as in Fig.1; dHb, dorsal habenula; vHb, ventral habenula. Scale bar=500μm in (b,h,n), 200μm in (c-d,i-j,o-p), 150μm in (e,k,q), 100μm in (f,l,r).

### Wnt signaling activity is dynamic and repressed on the left by Nodal in the catshark developing habenulae

To gain insight into the mechanisms controlling the formation of asymmetries shared by chondrichthyans, dipneusts, and polypterids in the lateral/ventral habenulae, we focused on the catshark, a model accessible to experimental approaches during development, and on canonical Wnt signaling. Expression of *ScLef1* and *ScTcf7l2*, which encode effectors of this pathway, was indeed characterized by statistically significant right enrichments in our transcriptomic analysis (q-value=2.2E-03 and 3.0E-03 respectively; Supplementary Table 2). We thus analyzed the dynamic of Wnt activity during habenula formation, by assessing the nuclear distribution of β-catenin on histological sections from the first appearance of habenular evaginations at stage 26 to stage 31 (Fig.3; Supplementary Fig.12). At stage 26, β-catenin is broadly distributed in the growing habenular evaginations but a nuclear signal is only observed in some HuC/D-positive cells, which form a discrete lateral cell population of differentiating neurons, transiently larger on the left than on the right side at this stage^27^ (Fig. 3a,b,a1-a4,b1-b4). A broad bilateral cytoplasmic signal is maintained at stage 28, albeit without clear nuclear restriction (Fig. 3c,c1-c4). A marked transition is observed at stage 29, with a nuclear accumulation of β-catenin restricted to the lateral right habenula and a withdrawal of the cytoplasmic signal in the lateral left one (Fig. 3d,d1-d4). This asymmetry is maintained at stage 31, with a high proportion of positive nuclei in the *Prox1-* expressing Right-LHb, while no signal (neither nuclear, nor cytoplasmic) is observed in the *Sox1*-positive Left-LHb (Fig. 3e, 3f; compare Fig.3e1-e2 with Fig.3e3-e4). The medial habenula (MHb, *Kctd12b*-positive; Supplementary Fig.12a,c,e) also expresses β-catenin at this stage but the signal is excluded from nuclei in all areas examined (Supplementary Fig.12b,d,f, b1-b2,d1-d8). However, a heterogeneity is observed within MHb, with an absence of cytoplasmic expression in its anterior left external component, known to express *ScPde1a* (Supplementary Fig.12b,d; compare Supplementary Fig.12d1-d2 to d3-d8). The restriction of Wnt signaling to the right habenula at stages 29-31 is reminiscent of the one reported in the zebrafish, as a result of a parapineal-mediated left repression^24,25^. In the absence of a parapineal in the catshark, we assessed the dependence of this asymmetry on Nodal signaling, by inhibiting the pathway during its window of left-restricted activity, shortly after neural tube closure as previously described^18^ (Fig. 3g,h). While control embryos exhibit the expected restriction of nuclear β-catenin to the lateral right habenula (Right-LHb; Fig. 3g,g1-g4), *in ovo* injection of the Nodal/Activin antagonist SB-505124 results in an expansion of the signal to the left one (Left-LHb), resulting in a right isomerism (Fig. 3h,h1-h4). Taken together, these data highlight an asymmetric, dynamic canonical Wnt activity in developing habenulae, and suggest that the marked restriction to the lateral right habenula observed starting from stage 29 is dependent on the transient left Nodal activity reported in the catshark diencephalon at earlier stages of development^18^.

**Figure 3.**
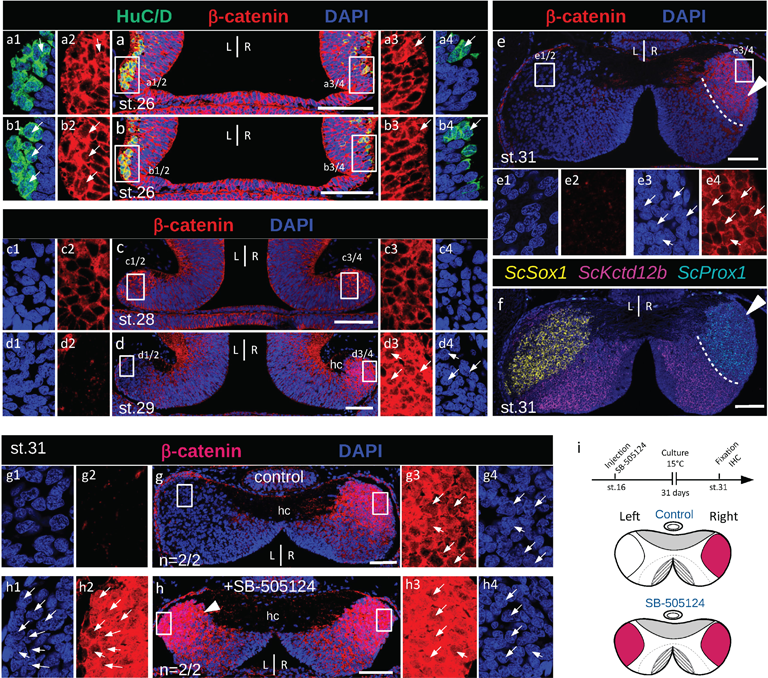
The nuclear β-catenin profile in developing catshark habenulae reveals dynamic, Nodal dependent asymmetries. **a-e** Confocal images of transverse sections of catshark habenulae at stages 26 (a,b), 28 (c), 29 (d) and 31 (e) following DAPI staining (blue) and immuno-histochemistry (IHC) with antibodies directed against β-catenin (red, a-e) and HuC/D (green, a-b). **f**Hybridization chain reaction-based fluorescent *in situ* hybridization (HCR-FISH) image of a section adjacent to (e) with territories for *ScSox1*, *ScKctd12b*, and *ScProx1* respectively in yellow, magenta, and cyan. White arrowheads in (e,f) point to Right-LHb. Dotted lines in (e,f) delimit Right-LHb and MHb subdomains, respectively positive for *ScProx1* and *ScKctd12b.* g-h Confocal images of transverse sections of stage 31 catshark habenulae, after *in ovo* injection of DMSO (g, control) or SB-505124 (h) following neural tube closure. The section shown in (a) is located anteriorly to the one shown in (b), sections in (c-h) are located at a medial organ level. A white arrowhead in (h) points to a lateral left territory showing a high density of β-catenin-positive nuclei, present in SB-505124-treated embryos but absent from control ones. **i** Schemes showing the experimental procedure and the habenular phenotypes observed, with lateral territories of nuclear β-catenin accumulation in red. Dorsal is to the top in all panels. Magnifications of boxed areas in (a-e,g-h) are shown in (a1-4 to e1-4,g1-4,h1-4). Thin white arrows point to β-catenin-labeled nuclei. Abbreviations: L, left; R, right. Scale bar=100μm.

### The elaboration of lateral right neuronal identities (Right-LHb) is dependent on Wnt signaling in the catshark

To test the involvement of the right-restricted Wnt activity detected in the lateral habenula in the elaboration of neuronal identities, we next inhibited the pathway using *in ovo* injection of the Wnt antagonist IWR-1, a tankyrase inhibitor that stabilizes the β-catenin degradation complex via Axin modification^30^ and analyzed the resulting phenotype at stage 31 (Fig.4, Supplementary Fig.13, Supplementary Table 3). A single injection of the drug at stage 29 results in a withdrawal of β-catenin nuclear signal from areas of varying size in the lateral right habenula (compare Fig.4a,a3,a4 and Fig.4g,g3,g4; Supplementary Fig.13a,e). Expression of the two Left-LHb markers *ScSox1* and *ScNtng2* expands to the right in all embryos injected with the drug (n=6/6; Supplementary Table 3), a phenotype never observed in uninjected or control embryos (compare Fig.4b,c,e and Fig.4h,i,k; Supplementary Fig.13b,f). This expansion is superimposable with the area devoid of nuclear β-catenin in the lateral right habenula (Figure 4g,h). Partial losses of two Right-LHb markers (*ScProx1* and *ScKiss1)* are consistently observed in the same subdomain (compare Fig.4d,f and Fig.4j,l; Supplementary Fig.13c,g). In the mouse, no asymmetric habenular expression of *Prox1* has been reported, but expression of the gene in adjacent thalamic territories has been shown to be dependent on *Tcf7l2*, in line with a dependence on Wnt signaling^31^. Another gene expressed in a neighboring thalamic region, *Rorα*, is submitted to the same regulation in this species. In view of the gene signature similarities between the catshark lateral habenula and mouse thalamic regions adjacent to the habenulae, we analyzed expression of the catshark *Rorα* ortholog in untreated, control and IWR-1 treated embryos. A lateral right-restricted expression territory of the gene, forming a radial band adjacent to the external part of MHb, is observed in habenulae of untreated embryos at stage 31 (Supplementary Fig.13d3), which is lost following IWR-1 treatment (compare Supplementary Fig.13d1,d2 with Supplementary Fig.13h1,h2). These data indicate that Wnt signaling is required for the elaboration of right neuronal identities and represses left ones in the catshark lateral right habenula (Fig. 4m). In contrast, we never observed changes of *ScEnpp2* and *ScPde1a* expressions in the medial habenula following IWR-1 treatment in these conditions (Suppl. Fig.14).

**Figure 4.**
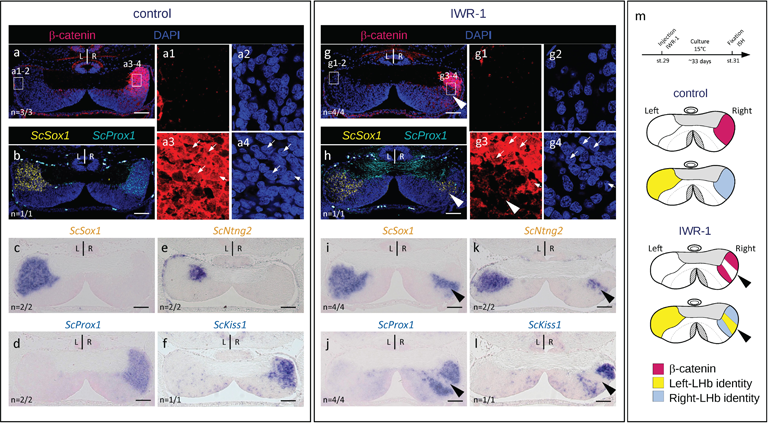
Inhibition of Wnt signaling converts lateral right into lateral left neuronal identities in developing catshark habenulae. **a-l** Transverse sections of catshark stage 31 habenulae from control (a-f) and IWR-1-treated (g-l) embryos, after IHC with an antibody directed against β-catenin (β-catenin in red and DAPI-stained nuclei in blue) (a,g), HCR-FISH with territories of *ScSox1* and *ScProx1* in yellow and cyan respectively (b,h), ISH with probes for *ScSox1* (c,i), *ScProx1* (d,j), *ScNtng2* (e,k) and *ScKiss1* (f,l). Sections are shown at a medial organ level, dorsal to the top. (a) and (b) show adjacent sections of the same embryo, same for (c) and (d), (e) and (f), (g) and (h), (i) and (j), (k) and (l). (a1-a4) and (g1-g4) show magnifications of the territories boxed in (a) and (g), with β-catenin signals in (a1,a3,g1,g3) and DAPI signals in (a2,a4,g2,g4). Thin white arrows point to β-catenin labeled nuclei. Arrowheads in (g-l) show expansions of Left-LHb markers, concomitant with the loss of Right-LHb markers and of nuclear β-catenin accumulation in the lateral right habenula. “n” in (a-l) refers to the number of embryos with the same phenotype for the markers and the detection method shown. **m** Schemes showing the experimental procedure used for pharmacological treatments and the resulting habenular phenotypes, with territories of nuclear β-catenin accumulation in red, territories of Left- and Right-LHb identity in yellow and blue respectively. Scale bar=100μm.

### The repression of Wnt signaling rescues lateral left neuronal identities in SB-505124 treated embryos

We previously showed that an early abrogation of Nodal/Activin signaling using in ovo injection of the inhibitor SB-505124 during the diencephalic window of left-restricted Nodal activity induces a right isomerism habenular phenotype^18^. To test whether Wnt inhibition could rescue lateral left (Left-LHb) neuronal identities in these embryos, we carried out double *in ovo* injections of SB-505124 (at stage 16) and IWR-1 (at stage 29) and analyzed the resulting phenotype at stage 31 (Fig.5; Supplementary Fig.15; Supplementary Table 4). As expected, *in ovo* injections of SB-505124 consistently results in a right isomerism (n=5/5), with a symmetric nuclear distribution of β-catenin in both the left and right lateral habenulae (Fig. 5a; Supplementary Fig.15a), a complete loss of lateral left expression for the two Left-LHb markers tested (*ScSox1* and *ScNtng2*: Fig.5c,d,f; Supplementary Fig.15b,c), and a concomitant lateral left expansion of Right-LHb markers (*ScProx1* and *ScKiss1*: Fig.5b,e,g; Supplementary Fig.15d). Following IWR-1 injection at stage 29 of SB-505124-treated embryos (n=7/7), the symmetry of habenulae is consistently maintained, but the lateral habenulae exhibit non-overlapping territories of Right- and Left-LHb identities, similar to the right habenulae of IWR-1-treated embryos. Compared to SB-505124-treated embryos, nuclear β-catenin is undetectable in lateral zones of variable size in both the left and the right habenulae (compare Fig.5h and 5a; Supplementary Fig.15a and 15e). Expression of Left-LHb markers (*ScSox1* and *ScNtng2*) is restored in the corresponding territories (compare Fig.5c,d,f and Fig.5j,k,m; Supplementary Fig.15b,c and 15f,g), while expression of Right-LHb markers (*ScProx1*, *ScKiss1*) is lost (compare Fig.5b,e,g and Fig.5i,l,n; Supplementary Fig.15b,d and 15f,h). Taken together, these data suggest that a repression of Wnt signaling is sufficient to promote a lateral left identity in the absence of Nodal signaling.

**Figure 5.**
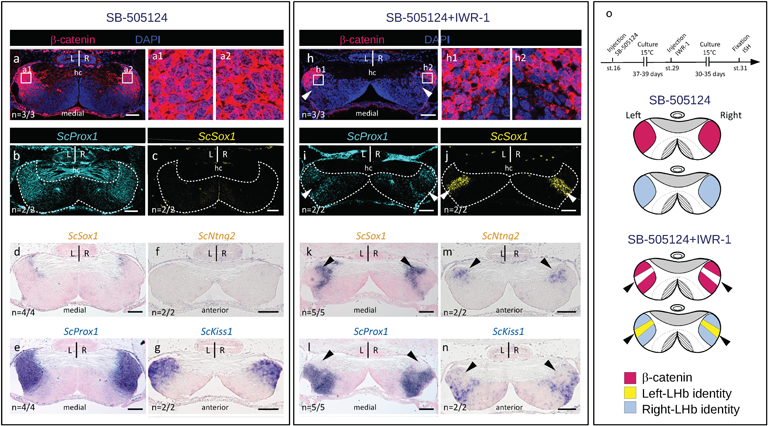
Inhibition of Wnt signaling rescues Left-LHb neuronal identities in the lateral habenulae of SB-505124 treated embryos. **a-n** Transverse sections of catshark stage 31 habenulae from SB-505124 (a-g) and SB-505124-IWR-1 treated (h-n) embryos, after IHC with an antibody directed against β-catenin (β-catenin in red, DAPI-stained nuclei in blue) (a,h), HCR-FISH with territories of *ScSox1* and *ScProx1* in yellow and cyan respectively (b,c,i,j), ISH with probes for *ScSox1* (d,k), *ScProx1* (e,l), *ScNtng2* (f,m) and *ScKiss1* (g,n). Sections are shown at anterior (f,g,m,n) or medial (a-e,h,n) organ levels, dorsal to the top. (a-c) show sections of the same embryo, same for (d,e), (f,g), (h-j), (k,l), and (m,n). (a1-a2) and (h1-h2) show magnifications of the territories boxed in (a) and (h), with β-catenin signals in red and DAPI signals in blue. A right isomerism is observed following SB-505124 treatment (a-g). Arrowheads in (h-n) point to bilateral LHb sub-territories harboring a Left-LHb identity as a result of the additional IWR-1 treatment. “n” in panels (a-n) refers to the number of embryos with the same phenotype for the markers and the detection method shown. **o** Schemes showing the experimental procedure used for pharmacological treatments and the resulting habenular phenotypes, with territories of nuclear β-catenin accumulation in red, territories of Left- and Right-LHb identity in yellow and blue respectively. Abbreviations: hc, habenular commissure; L, left; R, right. Scale bar=100μm.

### Left- and Right-LHb progenitors exit cell cycles prior to stage 29

In the zebrafish, Wnt signaling has been shown to delay neuronal differentiation in the dorsal habenulae, thus contributing to neuronal diversity and asymmetry^25^. In order to address whether the respective timing of progenitor cell cycle exits differs between the left and the right habenulae, we conducted BrdU (5-bromo-2’-deoxyuridine) pulse chase assays starting from early stages of neurogenesis^27^ (Fig.6; Supplementary Fig.16). BrdU incorporation pulses were carried out at stages 26-27 (two stages difficult to distinguish in live embryos), 28 and 29, and the distribution of BrdU-labeled cells was analyzed on transverse sections at different organ levels at stage 31. Assignment of labeled cells to the main habenular subdomains was achieved by comparisons to adjacent sections submitted to ISH (Fig. 6). On the right side, the maximal intensity of BrdU signals is observed along radial bands, which gradually regress from lateral to medial habenula levels at all stages analyzed (Fig. 6a2-a8,b2-b8,c2-c8). A negative territory restricted to posterior and lateral-most Right-LHb territories (*ScProx1-*positive and *ScKctd12b-*negative) is already present following a pulse at stages 26-27 (Fig. 6a4-a8), it expands after a pulse at stages 28 (compare Fig.6a4-a8 and Fig.6b1-b8), and the BrdU signal is completely lost in Right-LHb, becoming restricted to MHb, following a pulse at stage 29 (Fig. 6c1-c8). On the left side, a similar pattern is observed, with the presence of lateral posterior BrdU-negative territories after pulses at stages 26-27 (Fig. 6a7-a9) and a gradual withdrawal of BrdU-labeled cells from Left-LHb (*ScSox1*-positive and *ScKctd12b*-positive) after pulses between stages 28 and 29 (compare Fig.6b1-b9 with Fig.6c1-c9). However, a major difference is that on the left side, BrdU-negative territories span the anterior-most part of both the lateral (*ScSox1*-positive) and the medial (*ScKctd12b*-positive) habenulae following pulses at all stages analyzed, including stages 26-28 (Fig. 6a1-a6,b1-b6). Furthermore, the sharp boundary between BrdU-positive and −negative cells, observed on the right at all pulse stages and all habenula levels, is not observed on the left at anterior to medial organ levels and pulse stages preceding stage 28. Analyses of horizontal sections of stage 31 embryos submitted to BrdU pulses between stages 28 and 29 confirm these conclusions (Supplementary Fig.16a-b,e-f). BrdU-labeled cells are lost in both (*ScKctd12b* negative) Left- and Right-LHb following pulses between stage 28 and 29, regressing from external to internal MHb levels (Supplementary Fig.16c-d,e-f). These data indicate that at anterior-most organ levels, progenitors of both lateral and medial territories exit cell cycles earlier on the left side than on the right side (Fig. 6d,e).

**Figure 6.**
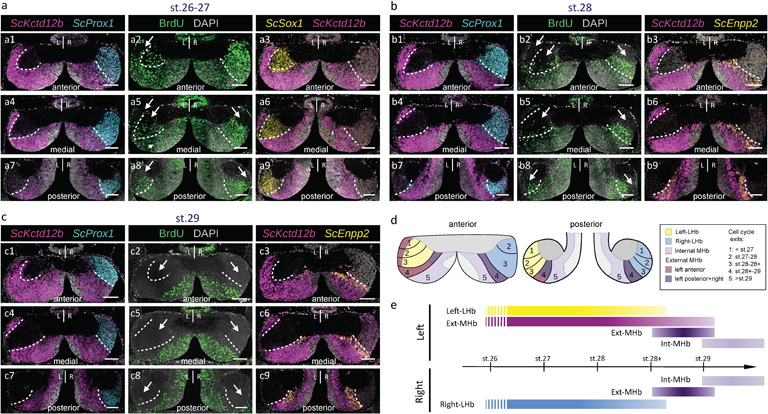
Spatial and temporal regulation of progenitor cell cycle exits in developing catshark habenulae. **a-d** Transverse sections of catshark stage 31 habenulae following exposure of embryos to BrdU pulses at stage 26-27 (a), 28 (b), and 29 (c), dorsal to the top. (a2,a5,a8,b2,b5,b8,c2,c5,c8) show confocal images following IHC using an antibody directed against BrdU (green), with DAPI-stained nuclei shown in gray. (a1,a4,a7,b1,b4,b7,c1,c4,c7), (a3,a6,a9), and (b3,b6,b9,c3,c6,c9) respectively show confocal images after double ISH with probes for *ScKctd12b* (magenta)/*ScProx1* (cyan), *ScSox1* (yellow)/*ScKctd12b* (magenta), and *ScKctd12b* (magenta)/*ScEnpp2* (yellow), with DAPI-stained nuclei in gray. (a1,a2,a3) show adjacent sections at an anterior level, same for (b1,b2,b3) and (c1,c2,c3). (a4,a5,a6) show adjacent sections at a medial level, same for (b4,b5,b6) and (c4,c5,c6). (a7,a8,a9) show adjacent sections at a posterior level, same for (b7,b8,b9) and (c7,c8,c9). White dotted lines delimit the border between medial (MHb) and lateral (LHb) territories as inferred from *ScKctd12b* expression, and its approximate location in BrdU-labeled sections. Thin arrows point to BrdU negative territories. **d** Scheme showing the spatial distribution of territories exiting the cell cycles earlier than stage 27 (1), between stages 27-28 (2), stages 28-28+ (3), stages 28+-29 (4), and later than stage 29 (5), superimposed on the subdomain organization of catshark stage 31 habenulae. **e** Scheme showing the developmental windows when the broad territories of stage 31 catshark habenulae exit the cell cycles. Data from Supplementary Figure 16 are taken into account in (e) and (f). Same color code in (d,e) as in Fig.1p. Scale bar=100μm.

### The distribution of nuclear β-catenin in lateral/ventral habenulae reflects asymmetry patterns in the lateral/ventral habenulae of gnathostomes

In the catshark, IHC profiles of β-catenin remain highly asymmetric in the habenulae of juveniles (Supplementary Fig.17), with nuclear signals restricted to the lateral right habenula (Supplementary Fig.17a7,a8,b3,b4,c5,c6). Heterogeneities are also observed in the medial habenula, with cytoplasmic signals being selectively absent in its external anterior left component (Supplementary Fig.17a3-a4). In order to test whether the presence of asymmetries in ventral/lateral habenulae correlates with asymmetric distributions of β-catenin across gnathostomes, we performed IHC analyses on transverse sections from reedfish and spotted gar juveniles, as well as frog tadpoles (stage NF-66) (Fig.7). The lungfish was excluded from this analysis as no positive control signal could be obtained in this species. In the reedfish (*E. calabaricus*), no nuclear signal could be observed on the left (Fig. 7a1,a2). In contrast, most of the right-restricted ventral territory of *Kiss1* expression contains a high density of β-catenin-positive nuclei (Fig.7a,a3,a4,7b;). In the dorsal right *EcKiss1-*negative territory, nuclei are generally unlabeled except in a limited lateral area (Fig. 7a5,a6). In contrast, in the spotted gar (*L. oculatus*), nuclear distribution of β-catenin is observed both on the right and on the left, in ventral *Kiss1*-positive territories (Fig. 7c,c1,c2,c3,c4,7d). As in the reedfish, dorsal *LoKiss1*-negative territories are heterogeneous, with some areas, such as the dorsal-most part of the left dorsal habenula, devoid of signal (Fig.7c), and others containing either cytoplasm-restricted or nuclear β-catenin signals (Fig. 7c5,c6). In the frog, no nuclear signals were observed either in the dorsal-most *Kctd*-positive or in the adjacent *Sox1*-positive domains (Fig. 5e,e1,e2,f,f1,f2,f3,f4). However, a significant proportion of β-catenin-positive nuclei is present in adjacent ventral territories, corresponding to those expressing *Prox1* (Fig. 7f5,f6,f7,f8). In conclusion, right-restricted nuclear accumulations of β-catenin, maintained until advanced stages of differentiation, correlate with the presence of neuronal identity asymmetries in ventral/lateral habenulae.

**Figure 7.**
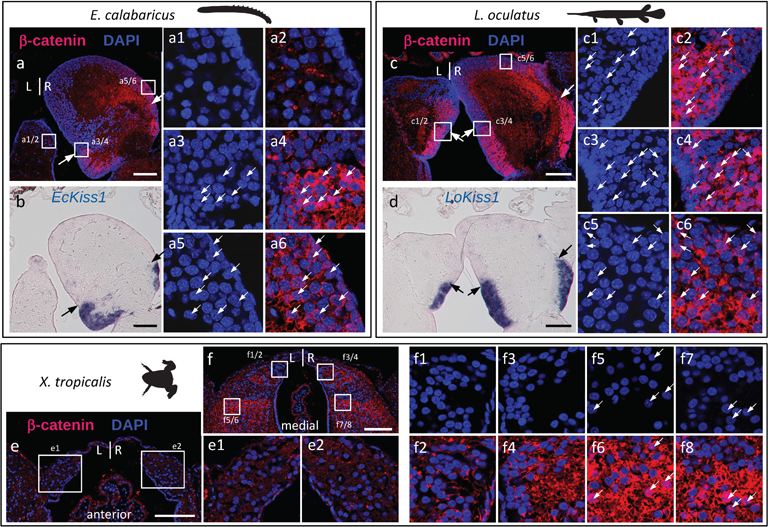
Evolution of nuclear β-catenin asymmetry patterns in the habenulae of jawed vertebrates. **a-f** Transverse sections of habenulae from reedfish (*Erpetoichthys calabaricus*; a-b), spotted gar (*Lepisosteus oculatus*; c-d), Western clawed frog (*Xenopus tropicalis*; e-f) juveniles after IHC using an antibody directed against β-catenin (red) with DAPI-stained nuclei in blue (a,c,e,f) and after ISH with probes for *Kiss1* orthologs (b,d). Dorsal is to the top in all panels. Arrows in (a,b,c,d) indicate the boundary between ventral, *Kiss1*-positive territories and dorsal territories. (a1-a6), (c1-c6), (e1-e2) and (f1-f8) show higher magnifications of territories boxed in (a), (c), (e) and (f) respectively, with white arrows pointing to β-catenin-positive nuclei. Scale bars=100μm.

## DISCUSSION

We show here that catshark habenulae exhibit a highly asymmetric organization, with a majority of asymmetric gene expression profiles reported for the first time in a vertebrate. Asymmetric profiles are detectable in both neural progenitors and neuronal territories, and they concern not only the medial (dorsal in teleosts) subdomain of habenulae as in the zebrafish, but also their lateral subdomain, which in the catshark contains very different neuronal identities between the left and the right side. This divergence between the zebrafish and the catshark questions the origin and the mode of evolution of habenular asymmetries across gnathostomes. Our phylogenetic approach reveals that, in the lateral/ventral subdomain of habenulae, orthologs of selected left- and right-restricted markers in the catshark share striking similarities in their asymmetry patterns, not only with the elephant shark, a member of holocephalans (which diverged from elasmobranchs about 410 million years ago^32^), but also with a sarcopterygian, the lungfish, and an actinopterygian, the reedfish. This asymmetry pattern contrasts with the bilateral expressions observed in lateral habenulae and adjacent ventral territories in the frog and the mouse, or in the ventral habenulae of the spotted gar and teleosts. While we cannot rule out convergent evolutionary processes to explain the similarities observed between chondrichthyans, dipneusts, and polypterids, we favor a more parsimonious hypothesis, and propose that the asymmetry pattern observed in catshark lateral habenulae reflects an ancestral trait of jawed vertebrates, independently lost in tetrapods and neopterygians (Fig. 8a). In support of this interpretation, the phylogenetic distribution of asymmetries, shared by chondrichthyans and members of the dipneusts and polypterids, respectively outgroups to tetrapods within sarcopterygians and to neopterygians within the actinopterygians, does not evoke a punctuated presence/absence pattern, as observed for instance for the left-restricted nucleus recently identified in teleosts^16^. Furthermore, the relative organization of habenular territories appears very similar in the mouse and the frog on the one hand, and within teleosts and in the spotted gar on the other hand^16^, arguing against a rapid drift of the corresponding molecular architectures in these taxa. Concerning the two tetrapods, it is intriguing that in the mouse, several orthologues of catshark Right-LHb markers (*Prox1*, *Rorα*, and *Rerg*) are not expressed in the habenulae but rather in the adjacent paraventricular nucleus of thalamus^31^. Similarly, in the frog, although the precise posterior and ventral boundaries of habenulae is difficult to identify due to the absence of marked morphological landmarks, we observed the same relative location of the *Prox1* territory, adjacent to *Sox1* expression in the lateral habenulae. Taken together, these data suggest that ancient regulatory circuits, controlling related neuronal identities, are deployed in different, habenular or thalamic, symmetric or asymmetric, territories depending on species.

**Figure 8.**
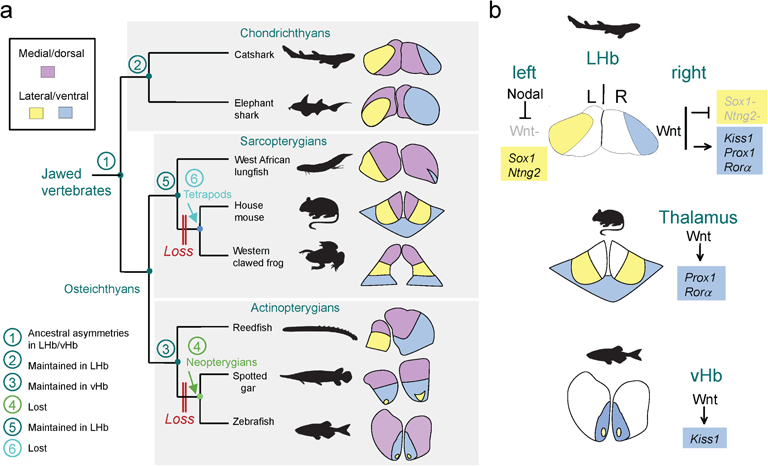
Evolution of habenular asymmetries in jawed vertebrates. **a** Distribution of neuronal identity asymmetries in lateral/ventral habenulae across jawed vertebrates. Schemes show the general subdomain organization of habenulae in eight members of the major gnathostome taxa, with medial/dorsal territories in purple, and lateral/ventral territories in yellow (territories of neuronal identity related to catshark Left-LHb) or blue (territories of neuronal identity related to catshark Right-LHb). Numbers at the nodes of the tree refer to the ancestral asymmetry profile inferred at this node, based on observations in members of the corresponding phyla. The phylogenetic distribution of asymmetries suggests that the neuronal identity asymmetries observed in the catshark lateral habenulae reflect an ancestral trait (node 1 of the tree), maintained in chondrichthyans (node 2) as well as in ancestral actinopterygians (node 3), persisting on the left side in ancestral sarcopterygians (node 5), but independently lost in neopterygians and tetrapods (nodes 4 and 6). **b** Differential deployment of an ancestral Wnt dependent regulatory module across jawed vertebrates. Together with data available in the mouse and the zebrafish, the functional analyses conducted in the catshark suggest that the Wnt dependence of *Kiss1/Prox1/Rora* expressions observed in the catshark lateral right habenula may reflect an ancient regulatory module, recruited to shape neuronal identities in symmetric territories (respectively ventral habenula and dorsal thalamus) of neopterygians and tetrapods.

What could be the underlying mechanism? In the catshark, our data highlight a role for Wnt signaling in the elaboration of lateral right versus left neuronal identities (Fig. 8b). Contrary to the involvement of Wnt signaling in the elaboration of asymmetries in the zebrafish dorsal habenula, the conversion of Right-into Left-LHb neuronal identities observed in the lateral right habenula following Wnt inactivation is unlikely to result from perturbations of the temporal regulation of neurogenesis^25,33^. Our BrdU analyses rather suggest that Left- and right-LHb progenitors exit the cell cycles largely simultaneously, and that Right- and Left-LHb precursors were postmitotic at the stage when Wnt signaling was inactivated. This is, however, reminiscent of the role of *Tcf7l2*, an effector of Wnt signaling required for the refinement of neuronal identities in thalamic nuclei adjacent to the habenula and for the expression of *Prox1* and *Rorα* in postmitotic neurons in the mouse^31,34^. Similarly, the zebrafish orthologue of *Tcf7l2* is required for the specification of *kiss1* expression in the ventral habenula, suggesting a conserved Wnt-dependent regulation of this gene between the zebrafish-symmetric-ventral habenulae and the catshark lateral right habenula^23^. Finally, the detection of nuclear β-catenin in the ventral habenulae of the spotted gar, in the right ventral habenulae of the reedfish and in the *Prox1* expressing region adjacent to the lateral habenulae of the xenopus is consistent with a deep conservation of Wnt signaling functions in the elaboration of neuronal identities, even though the cellular contexts vary between species. We therefore propose that the Wnt-dependence of lateral right neuronal identities observed in the catshark reflects an ancient regulation, which controlled asymmetry formation in the lateral habenulae of ancestral jawed vertebrates and which has been deployed in different, bilateral territories of tetrapods and neopterygians.

How these mechanistic transitions have taken place remains elusive. Despite the different nature of asymmetries in the catshark lateral habenulae and the zebrafish dorsal habenulae, our analysis highlights many similarities in their mechanisms of formation. In both cases, a left-sided Wnt repression is essential, even though it is dependent on Nodal signaling in the catshark and on an interaction with the parapineal in the zebrafish^24^. Whether the left restricted expression of *Sox1* in the catshark habenula, may be involved in this repression, similar to the role of *sox1a* in the zebrafish parapineal^35^, is an intriguing possibility. We also find that Wnt activity is submitted to a highly dynamic regulation in the catshark as in the zebrafish^22,25^. Finally, neurogenesis in the developing habenulae follows a strict temporal pattern in the catshark, resulting in the characteristic lateral to medial organization in radial bands observed. A similar temporal regulation of neuronal identities, connectivity patterns and functional properties, has also been described in the mouse and the zebrafish habenulae^7,33,36^. The asymmetry of this process in the zebrafish results in an advanced differentiation of dorsal habenula neurons on the left compared to the right^25^, which is reminiscent of the early differentiation of MHb neurons on the left in the catshark. Comparisons between the catshark and the mouse suggest that related neuronal identities might be specified under the control of this sequential diffferentiation process, as two genes specifically expressed in the lateral-most territory of the mouse medial habenulae (*Spon1* and *Trhde*) have a similar expression territory in the early-born left subdomain of the medial habenula (external anterior left, *Pde1a*-positive subdomain) in the catshark. The similarities observed in the lateral habenulae between the catshark left side and the mouse (both left and right sides) might therefore be extended to some zones of the medial habenulae, raising the question of an underlying mechanistic conservation. A possible candidate for a conserved actor in this regulation is Wnt signaling, as it is responsible for the delayed differentiation of dorsal habenula neurons on the right relative to the left in the zebrafish^25,33^. In line with this hypothesis, we observed differences in cytoplasmic β-catenin profiles between left-restricted, early born, and bilateral, later born catshark MHb territories, suggesting a relationship between the timing of neurogenesis and the regulation of Wnt pathway components.

Taken together, our data support the existence of a conserved regulatory logic for the formation of habenular asymmetries across gnathostomes, with a coupling between a conserved temporal regulation of neurogenesis, and its modification by a repression of Wnt signaling, restricted to the left in the ancestral state. Due to its highly dynamic regulation, the latter may be prone to gradual variations in time and space, allowing the quick diversification of asymmetry developmental processes. Such variations may in turn lay the ground for major mechanistic transitions, such as the switch between a Nodal-dependent left repression of Wnt signaling, as observed in the lamprey and catshark, to a parapineal dependent repression, as described in the zebrafish^18^.

## METHODS

### RNA isolation

Left and right habenulae were manually dissected from a total of 45 anaesthetized stage 31 catshark embryos and stored in RNAlater (Thermo Fisher Scientific, Carlsbad, CA, USA) until RNA extraction. Three left pools, each containing 15 left habenulae and three right pools, containing the corresponding 15 right habenulae, were prepared from these explants. Total RNA was extracted from these pools using the Ribopure Kit (Thermo Fisher Scientific, Carlsbad, CA, USA). RNA quantities and integrity indexes (RINs) were controlled using a Bioanalyzer 2100 (Agilent Technologies, Santa Clara, CA, USA). RNA quantities ranging between 1.1 and 2.4 µg and RIN values comprised between 7.4 and 9.6 were obtained for the analyzed pools.

### Library construction and sequencing

We prepared RNA-seq libraries using the Truseq Stranded mRNA Sample Prep Kit (20040532, Illumina, San Diego, CA, USA) according to the manufacturer’s instructions. Briefly, polyadenylated RNAs were selected using oligo-dT magnetic beads. The polyA+ RNAs were fragmented using divalent cations at elevated temperature and reverse transcribed using random hexamers, Super Script II (18064014, Thermo Fisher Scientific, Carlsbad, CA, USA) and actinomycin D. Deoxy-TTP was replaced by dUTP during the second strand synthesis to prevent its amplification by PCR. Double stranded cDNAs were adenylated at their 3’ ends and ligated to Illumina’s adapters containing unique dual indexes (UDI). Ligated cDNAs were PCR amplified in 15 cycles, and the PCR products were purified using AMPure XP Beads (A63880, Beckman Coulter Genomics, Brea, CA, USA). The size distribution of the resulting libraries was monitored using a Fragment Analyzer (Agilent Technologies, Santa Clara, CA, USA) and the libraries were quantified using the KAPA Library Quantification Kit (Roche, Basel, Switzerland). The libraries were denatured with NaOH, neutralized with Tris-HCl, and diluted to 7.5 pM. Clustering was carried out on a cBot and sequencing was performed on a HiSeq 2500 (Illumina, San Diego, CA, USA) using the paired-end 2×125 nt protocol on two lanes of a flow cell (SRA: PRJNA1040760).

### Mapping and expression profiling

Reads obtained from each one of the three left and right habenula pools were pseudo-mapped onto an annotated database of reference gene models^37^, and pseudo-counted using a k-mer quantification method, kallisto^38^. Contigs exhibiting statistically significant count differences between the left and right habenulae were identified using the Wald test (q-value threshold 5E-02) implemented in sleuth^39^.

### Gene ontology (GO) analysis

Left versus right habenulae GO term enrichment gene analysis was carried out using the ConsensusPathDB online tool (http://cpdb.molgen.mpg.de/)^40,41^ against the whole list of genes in the catshark gene model reference. It was restricted to levels 2-5 GO terms related to biological processes (p-value cut-off 5E-02, further curated to a q-value threshold of 5E-02).

### Embryo collection

*Scyliorhinus canicula* eggs were provided by the Aquariology Service of the Banyuls-sur-Mer Oceanological Observatory. Embryos were dissected, fixed, and staged as previously described^42^. Juvenile of *Lepisosteus oculatus* and *Erpetoichthys calabaricus* were provided from commercial sources (https://www.poisson-or.com/). Juveniles of *Callorhincus milii* were provided by C. Boisvert (Curtin University, Australia) and *Xenopus tropicalis* by S. Marcellini (University of Concepcion, Chile) and N. Pollet (Université Paris-Saclay, France).

### *In situ* hybridization (ISH), hybridization chain-reaction based fluorescent in situ hybridization (HCR-FISH), and immunohistochemistry (IHC)

Probes were obtained from collections of embryonic *S. canicula* cDNA recombinants^43^, or obtained from synthetic double-stranded DNA (Suppl. Table 5). Following ISH, nuclei were counterstained using Nuclear Fast Red Solution (N3020, Sigma-Aldrich, Saint-Louis, MO, USA). Chromogenic ISHs of paraffin sections were carried out using digoxigenin-labeled antisense RNAs and standard protocols as previously described. Fluorescent ISHs were performed using digoxigenin- and fluorescein-labeled antisense RNAs following a published protocol^44^. HCR-FISH were carried out using published methods^45,46^ with slight modifications: slides were incubated for 20 min in 10µg/mL proteinase K (RPROTK-RO, Sigma-Aldrich, Saint-Louis, MO, USA) at room temperaturebefore probe hybridization. For each gene, probes were designed by Molecular Instrument (Los Angeles, CA, USA) using the following NCBI accession numbers: XM_038775015.1 (*ScKctd12b*), XM_038811423.1 (*ScProx1*) and XMsing the following NCBI accession numbers XM_038775015.1 (*ScKctd12b*), XM_038811423.1 (*ScProx1*) and XM_038818902.1 (*ScSox1*). The following amplifiers were used for the respective probes: B3-Alexa Fluor594, B32-Alexa Fluor488 and B1-Alexa Fluor647 were Amplifier respectively used. Fluorescent IHC of β-catenin (Ab6302, 1/1000, Abcam, Boston, MA, USA) on sections was carried out as previously described^27^. For fluorescent ISHs, HCRs, and IHCs, sections were imaged on a Leica SP8 confocal laser-scanning microscope.

### BrdU pulse chase analysis

BrdU (5-bromo-2’-deoxyuridine) pulse labelling and BrdU detection were performed as previously described^27^ with the following modifications: catshark embryos were removed from the egg case, incubated in oxygenated filtered sea water containing 5 mg/ml BrdU (16.2mM) (B5002, Sigma-Aldrich, Saint-Louis, MO, USA) for 16 hours and, for the chase, transferred to filtered sea water at 16°C until the desired stages. Following incubation of sections with the anti-BrdU primary antibody (sc-32323, 1/100, Santa Cruz Biotechnology, Dallas, TX, USA), detection was carried out with the mouse IgG kappa binding protein conjugated to horseradish peroxidase (1/100) (sc-516102, Santa Cruz Biotechnology, Dallas, TX, USA) and the TSA Plus Cyanine 3 Kit (NEL744001KT, Akoya Biosciences, Menlo Park, CA, USA) following the supplier’s instructions.

### Pharmacological treatments

For Nodal/Activin signaling inhibition, catshark embryos were treated at stage 16 by *in ovo* injection of 100 μl of a 50 µM solution of SB-505124 (S4696, Sigma-Aldrich, Saint Louis, MO, USA, a selective inhibitor of TGF-β type I receptors Alk4/5/7/18, as previously described ^18^. For Wnt inactivation, 100 μl of a DMSO-based solution containing 1 mM IWR-1 (I0161, Sigma-Aldrich, Saint-Louis, MO, USA), a selective inhibitor of tankyrase known to inhibit the cytoplasmic accumulation of β-catenin through stabilization of the destruction complex member Axin2, were injected into the egg case at stage 29, and eggs were maintained in oxygenated sea water at 16°C until stage 31. The same protocol was applied to control embryos, except for the absence of the drugs in the injected solutions.

## ACKNOWLEDGEMENTS

We thank Pascal Romans and the Centre de Ressources Biologiques Marines the Banyuls Oceanological Observatory (OOB) for help in obtaining specimens, the Sanger Institure for providing access to the catshark genome prior to public release, EMBRC-France for support to local marine infrastructures, David Pecqueur and the BioPic imaging platform for access to confocal microscopy, the Bio2Mar service for access to molecular biology platform, the UMR7232 Service de Bio-informatique BSBII for HM’s support.

## AUTHOR CONTRIBUTIONS

SM (Sylvain Marcellini), NP, EC, MDT, CB, MS, PB, and SM (S. Mazan) conceived the study and its experimental design in the different model organisms analyzed; ML, LM, RL, LG, VL, DS, KM, HC performed the experiments; HM, CK, and BB performed bioinformatic analyses; SM (S. Marcellini), MDT, MS and SM (S. Mazan) wrote the manuscript. All authors read and approved the manuscript.

## REFERENCES

1. Matsumoto, M. & Hikosaka, O. Lateral habenula as a source of negative reward signals in dopamine neurons. Nature 447, 1111–1115 (2007).

2. Stamatakis, A. M. & Stuber, G. D. Activation of lateral habenula inputs to the ventral midbrain promotes behavioral avoidance. Nat Neurosci 15, 1105–1107 (2012).

3. Beretta, C. A., Dross, N., Guiterrez-Triana, J. A., Ryu, S. & Carl, M. Habenula circuit development: Past, present, and future. Frontiers in Neuroscience 6, 1–10 (2012).

4. Andalman, A. S. et al. Neuronal Dynamics Regulating Brain and Behavioral State Transitions. Cell 177, 970–985.e20 (2019).

5. Amo, R. et al. Identification of the zebrafish ventral habenula as a homolog of the mammalian lateral habenula. The Journal of neuroscience: the official journal of the Society for Neuroscience 30, 1566–1574 (2010).

6. Hashikawa, Y. et al. Transcriptional and Spatial Resolution of Cell Types in the Mammalian Habenula. Neuron 106, 743–758.e5 (2020).

7. van de Haar, L. L. et al. Molecular signatures and cellular diversity during mouse habenula development. Cell Reports 40, 111029 (2022).

8. Pandey, S., Shekhar, K., Regev, A. & Schier, A. F. Comprehensive Identification and Spatial Mapping of Habenular Neuronal Types Using Single-Cell RNA-Seq. Current Biology 28, 1052–1065.e7 (2018).

9. Ahumada-Galleguillos, P. et al. Directional asymmetry in the volume of the human habenula. Brain Struct Funct. 222, 1087–1092 (2016).

10. Concha, M. L. & Wilson, S. W. Asymmetry in the epithalamus of vertebrates. Journal of Anatomy 199, 63–84 (2001).

11. Roussigné, M., Blader, P. & Wilson, S. W. Breaking symmetry: The zebrafish as a model for understanding left-right asymmetry in the developing brain. Developmental Neurobiology 72, 269–281 (2012).

12. Facchin, L., Duboué, E. R. & Halpern, M. E. Disruption of Epithalamic Left–Right Asymmetry Increases Anxiety in Zebrafish. J. Neurosci. 35, 15847–15859 (2015).

13. Chen, W. et al. Role of Olfactorily Responsive Neurons in the Right Dorsal Habenula–Ventral Interpeduncular Nucleus Pathway in Food-Seeking Behaviors of Larval Zebrafish. Neuroscience 404, 259–267 (2019).

14. Dreosti, E., Vendrell Llopis, N., Carl, M., Yaksi, E. & Wilson, S. W. Left-right asymmetry is required for the habenulae to respond to both visual and olfactory stimuli. Current Biology 24, 440–445 (2014).

15. Duboué, E. R., Hong, E., Eldred, K. C. & Halpern, M. E. Left Habenular Activity Attenuates Fear Responses in Larval Zebrafish. Current Biology 27, 2154–2162.e3 (2017).

16. Michel, L. et al. Diversification of habenular organization and asymmetries in teleosts: Insights from the Atlantic salmon and European eel. Front. Cell Dev. Biol. 10, 1015074 (2022).

17. Villalón, A. et al. Evolutionary plasticity of habenular asymmetry with a conserved efferent connectivity pattern. PLoS ONE 7, e35329 (2012).

18. Lagadec, R. et al. The ancestral role of nodal signalling in breaking L/R symmetry in the vertebrate forebrain. Nature communications 6, 6686 (2015).

19. Gamse, J. T., Thisse, C., Thisse, B. & Halpern, M. E. The parapineal mediates left-right asymmetry in the zebrafish diencephalon. Development 130, 1059–1068 (2003).

20. Gamse, J. T. et al. Directional asymmetry of the zebrafish epithalamus guides dorsoventral innervation of the midbrain target. Development 132, 4869–4881 (2005).

21. deCarvalho, T. N., et al. Neurtransmitter map of the asymmetric dorsal habenular nuclei of Zebrafish. Genesis 52, 636–655 (2014).

22. Kuan, Y.-S. et al. Distinct requirements for Wntless in habenular development. Developmental Biology 406, 117–128 (2015).

23. Beretta, C. A., Dross, N., Bankhead, P. & Carl, M. The ventral habenulae of zebrafish develop in prosomere 2 dependent on Tcf7l2 function. Neural development 8, 19 (2013).

24. Hüsken, U. et al. Tcf7l2 is required for left-right asymmetric differentiation of habenular neurons. Current Biology 24, 2217–2227 (2014).

25. Guglielmi, L. et al. Temporal control of Wnt signaling is required for habenular neuron diversity and brain asymmetry. Development 147, dev182865 (2020).

26. Coolen, M. et al. The dogfish Scyliorhinus canicula: A reference in jawed vertebrates. Cold Spring Harbor Protocols 3, 1–14 (2008).

27. Lagadec, R. et al. Neurogenetic asymmetries in the catshark developing habenulae: mechanistic and evolutionary implications. Sci Rep 8, 4616 (2018).

28. Lein, E. S. et al. Genome-wide atlas of gene expression in the adult mouse brain. Nature 445, 168–176 (2007).

29. Kan, L. et al. Dual function of Sox1 in telencephalic progenitor cells. Developmental Biology 310, 85–98 (2007).

30. Huang, S.-M. A. et al. Tankyrase inhibition stabilizes axin and antagonizes Wnt signalling. Nature 461, 614–620 (2009).

31. Lipiec, M. A. et al. TCF7L2 regulates postmitotic differentiation programs and excitability patterns in the thalamus. Development dev.190181 (2020) doi:10.1242/dev.190181.

32. Renz, A. J., Meyer, A. & Kuraku, S. Revealing Less Derived Nature of Cartilaginous Fish Genomes with Their Evolutionary Time Scale Inferred with Nuclear Genes. PLoS ONE 8, e66400 (2013).

33. Aizawa, H., Goto, M., Sato, T. & Okamoto, H. Temporally Regulated Asymmetric Neurogenesis Causes Left-Right Difference in the Zebrafish Habenular Structures. Developmental Cell 12, 87–98 (2007).

34. Lee, M. et al. Tcf7l2 plays crucial roles in forebrain development through regulation of thalamic and habenular neuron identity and connectivity. Developmental Biology 424, 62–76 (2017).

35. Lekk, I. et al. Sox1a mediates the ability of the parapineal to impart habenular left-right asymmetry. eLife 8, e47376 (2019).

36. Fore, S. et al. Functional properties of habenular neurons are determined by developmental stage and sequential neurogenesis. Sci. Adv. 6, eaaz3173 (2020).

37. Mayeur, H. et al. When Bigger Is Better: 3D RNA Profiling of the Developing Head in the Catshark Scyliorhinus canicula. Front. Cell Dev. Biol. 9, 744982 (2021).

38. Bray, N. L., Pimentel, H., Melsted, P. & Pachter, L. Near-optimal probabilistic RNA-seq quantification. Nat Biotechnol 34, 525–527 (2016).

39. Pimentel, H., Bray, N. L., Puente, S., Melsted, P. & Pachter, L. Differential analysis of RNA-seq incorporating quantification uncertainty. Nat Methods 14, 687–690 (2017).

40. Kamburov, A., Wierling, C., Lehrach, H. & Herwig, R. ConsensusPathDB—a database for integrating human functional interaction networks. Nucleic Acids Research 37, D623–D628 (2009).

41. Kamburov, A. et al. ConsensusPathDB: toward a more complete picture of cell biology. Nucleic Acids Research 39, D712–D717 (2011).

42. Ballard, W. W., Mellinger, J. & Lechenault, H. A series of normal stages for development of Scyliorhinus canicula, the lesser spotted dogfish Chondrichthyes: Scyliorhinidae. Journal of Experimental Zoology 267, (1993).

43. Coolen, M. et al. Evolution of axis specification mechanisms in Jawed vertebrates: Insights from a chondrichthyan. PLoS ONE 2, e374 (2007).

44. Lauter, G., Söll, I. & Hauptmann, G. Sensitive Whole-Mount Fluorescent In Situ Hybridization in Zebrafish Using Enhanced Tyramide Signal Amplification. in Brain Development (ed. Sprecher, S. G.) vol. 1082 175–185 (Humana Press, 2014).

45. Choi, H. M. T. et al. Third-generation in situ hybridization chain reaction: Multiplexed, quantitative, sensitive, versatile, robust. Development (Cambridge) 145, (2018).

46. Schwarzkopf, M. et al. Hybridization chain reaction enables a unified approach to multiplexed, quantitative, high-resolution immunohistochemistry and *in situ* hybridization. Development 148, dev199847 (2021).

