## Supplementary Files for "Analysis of a shark reveals ancient, Wnt dependent, habenular asymmetries in jawed vertebrates"

### SUPPLEMENTARY MATERIALS

**Supplementary Figure 1. Functional annotation of left- and right-enriched genes identified by transcriptomic analysis. a** Scheme showing the pipeline used for the functional annotation of genes identified by the transcriptomic comparison between left and right catshark habenulae. **b-d** Lists of over-represented GO- (Gene Ontology) terms retrieved from both left- and right-enriched gene lists (b), only from the list of left-enriched genes (c) or only from the list of right-enriched genes (d). For each term, the corresponding p-value (shaded in yellow for left-enriched genes and in blue for right-enriched genes) is indicated. Abbreviations: GPCR, G protein coupled receptor; L, left; R, right.

**Supplementary Figure 2. Expression profiles of left- or right-enriched genes showing regionalized ISH profiles in stage 31 catshark habenulae**. **a** Schemes showing a left lateral view of a stage 31 embryonic brain (a1; habenulae boxed in red), a lateral left view of the habenulae with a dotted line indicating the transverse section plane used (a2), and the subdomain organization observed on a section at a medial level. A dotted line delimits the external and internal components of the medial habenula. Color code: yellow, Left-LHB; light purple, MHb; blue, Right-LHB; magenta, anterior left external MHb; dark purple, right plus posterior left external MHb; hatched, proliferative pseudo-stratified neuroepithelium. **b-h** transverse sections following ISH with probes for Left-LHb markers (b), anterior-left external MHb markers (c), MHb markers (d), right+posterior-left external MHb markers (e), Right-LHb markers (f), genes expressed in combinations of LHb and MHb territories (g) and neural progenitor (LVZ) markers (h). Dorsal is to the top for all sections. For each gene, the rank (r), based on q-value in the list of asymmetrically expressed genes, and the laterality of the enrichment are indicated at the bottom left of the panel. Black and white arrowheads in (b,f) respectively point to Left- and Right-LHb labeled territories, thin arrows in (c,e) point to the boundary between complementary anterior and posterior territories within the left external MHb. Abbreviations: ant., anterior; post., posterior; LVZ, lateral ventricular zone; LHb, lateral habenula; MHb, medial habenula; hc, habenular commissure; L, left; R, right; pi, pineal. Scale bar=100µm.

**Supplementary Figure 3. Expression of *ScSox1*, *ScPde1a*, *ScKctd12b*, *ScEnpp2* and *ScProx1* along the antero-posterior axis of stage 31 catshark habenulae. a** Schemes showing a left lateral view of stage 31 catshark habenulae, with transverse section planes indicated by dotted lines (a1) and the subdomain organization observed on a section at a medial level of the structure (a2). Color code: yellow, Left-LHB; light purple, internal MHb; blue, Right-LHB; magenta, anterior left external MHb; dark purple, right plus posterior left external MHb; hatched, proliferative pseudo-stratified neuroepithelium. **b-f** Transverse sections after ISH with probes for *ScSox1* (b), *ScPde1a* (c), *ScKctd12b* (d), *ScEnpp2* (e) and *ScProx1* (f), dorsal to the top of each panel. All sections shown were obtained from the same embryo. (b1-f1), (b2-f2), (b3-f3), (b4-f4) and (b5-f5) each show successive adjacent sections. (b1-5), (c1-5), (d1-5), (e1-5) and (f1-5) each show sections from anterior to posterior levels as indicated in (a2). Arrowheads point to lateral habenulae, with black and white arrowheads respectively for the Left- and Right-LHb. Asterisks in (b5) and (f4) show minor *ScSox1* and *ScProx1* posterior expression sites, contra-lateral to their main lateral territory. Dotted lines delimit the subdivision of the medial habenula into external and internal components, as inferred from inner boundaries of the *ScPde1a* and *ScEnpp2* territories. Thin black arrows indicate the boundary between the complementary *ScPde1a* and *ScEnpp2* territories within the left external medial habenula. Abbreviations: hc, habenular commissure, pi, pineal; LHb lateral habenula; MHb, medial habenula; ext., external; int., internal, L, left;R, right. Scale bar=100µm.

**Supplementary Figure 4. Expression of *ScSox1*, *ScPde1a*, *ScKctd12b*, *ScEnpp2* and *ScProx1* along the dorso-ventral axis of stage 31 catshark habenulae. a** Schemes showing a left lateral view of stage 31 catshark habenulae, with horizontal section planes indicated by dotted lines (a1) and the subdomain organization observed on a section at a medial level of the structure (a2). Same color code as in Supplementary Fig.3. **b-f** Horizontal sections after ISH with probes for *ScSox1* (b), *ScPde1a* (c), *ScKctd12b* (d), *ScEnpp2* (e) and *ScProx1* (f), anterior to the top of each panel. All sections were obtained from the same embryo. (b1-f1), (b2-f2), (b3-f3) and (b4-f4) each show successive adjacent sections. (b1-4), (c1-4), (d1-4), (e1-4) and (f1-4) each show sections from dorsal to ventral levels as indicated in (a1). Arrowheads in (b,f) point to the lateral habenulae, with black and white arrowheads respectively for Left- and Right-LHb. Asterisks in (b4) and (f3) show minor *ScSox1* and *ScProx1* posterior expression sites, contra-lateral to their main lateral territory. Dotted lines delimit the subdivision of the medial habenula into external and internal components, as inferred from inner boundaries of the *ScPde1a* and *ScEnpp2* territories. Thin black arrows indicate the boundary between the complementary *ScPde1a* and *ScEnpp2* territories within the left external medial habenula. Same abbreviations as in Supplementary Fig.3. Scale bar=100µm.

**Supplementary Figure 5. Subdomain organization of juvenile catshark habenulae. a** Schemes showing a lateral view of juvenile catshark habenulae, with section planes used in (b-k) indicated by dotted lines (a1), and the subdomain organization observed on a transverse section at an anterior level of the structure (a2). Same color code as in Supplementary Fig.3. **b-f** Transverse sections after ISH with probes for *ScSox1* (b), *ScPde1a* (c), *ScKctd12b* (d), *ScEnpp2* (e) and *ScProx1* (f), dorsal to the top of each panel. Sectiions (b-f) were obtained from the same specimen. (b1-f1), (b2-f2), (b3-f3) and (b4-f4) each show successive adjacent sections. (b1-4), (c1-4), (d1-4), (e1-4), and (f1-4) each show sections from anterior to posterior levels as indicated in (a1). **g-k** Horizontal sections at a medial level of habenulae after ISH with probes for *ScSox1* (g), *ScPde1a* (h), *ScKctd12b* (i), *ScEnpp2* (j) and *ScProx1* (k), anterior to the top of each panel. Sections (g-k) were obtained from the same specimen. (g), (h), (i), (j) and (k) show successive adjacent sections at the levels indicated in (a1). **l-p** Transverse sections after ISH with probes for *ScNtng2* (l), *ScPcdh17* (m), *ScKctd12a* (n), *ScKctd8* (o) and *ScKiss1* (p), dorsal to the top of each panel. Arrowheads point to the lateral habenulae, with black and white arrowheads respectively for Left- and Right-LHb. Asterisks in (b1,b2,f3) show minor *ScSox1* and *ScProx1* posterior xpression sites, contra-lateral to their main lateral territory. Dotted lines delimit the subdivision of the medial habenula into external and internal components, as inferred from inner boundaries of the *ScPde1a* and *ScEnpp2* territories. Thin black arrows indicate the boundary between complementary anterior (*ScPde1a*-positive) and posterior (*ScEnpp2*-positive) subdomains of the left external medial habenula. Same abbreviations as in Supplementary Fig.3. Scale bar=100µm.

**Supplementary Figure 6. Expression of mouse orthologues of catshark Left-LHb, MHb and Right-LHb markers. a-k** Images of mouse adult coronal (a-i) or sagittal (j-k) sections following ISH with probes for *Kctd8* (a), *Spon1* (b), *Trhde* (c), *Ntng2* (d), *Pcdh17* (e), *Prkcq* (f), *Rora* (g), *Prox1* (h), *Stxbp6* (i), *Rerg* (j) and *Pde1a* (k) at the level of habenulae. All images were taken from the Allen Brain Atlas. Gene names are shown with the following color code: orthologues of markers of catshark Left-LHb, yellow; external anterior Left-MHb, magenta; MHb, dark purple; Right-LHb, blue. Abbreviations: LHb, lateral habenula; lMHb, lateral medial habenula; vMHb, ventral medial habenula; MHb, medial habenula; PVT, paraventricular nucleus of the thalamus. Scale bar=420µm in (a-i) and 840µm in (j-k).

**Supplementary Figure 7. Subdomain organization of habenulae in the elephant shark *Callorhinchus milii*. a** Schemes showing a left lateral view of the elephant shark brain, with the location of habenulae in red and dotted lines indicating transverse section planes at anterior, medial and posterior levels (a1), and a section at a posterior organ level with the habenular commissure in gray (a2). **b-i** Transverse sections of elephant shark juvenile habenulae after ISH with probes for

*CmPcdh17* (b), *CmNtng2* (c), *CmSox1* (d), *CmKctd8* (e), *CmKctd12a* (f), *CmKctd12b* (g), *CmKiss1* (h), and *CmProx1* (i), dorsal to the top. All sections were obtained from the same specimen. The level of the sections along the antero-posterior axis is indicated on each panel. Black and white arrowheads point to lateral territories respectively co-expressing *CmPcdh17*/*CmNtng2*/*CmSox1* on the left, and *CmKiss1/CmProx1* on the right. Thin arrows indicate their boundary with *CmKctd8/12a/12b*-positive, medial habenula territories. Asterisks in (b,h1,h2) indicate medial territories where *CmPcdh17* (b) and *CmKiss1* (h1,h2) are expressed in addition to their major lateral territories. Dotted lines in (e,f,g) delimit two subdomains within the left medial habenula, respectively positive or negative for *Kctd12a*. Abbreviations: hc, habenular commissure; Mes, mesencephalon; Tel, telencephalon; L, left; R, right. Scale bar=500µm.

**Supplementary Figure 8. Subdomain organization of habenulae in the reedfish *Erpetoichthys calabaricus*. a** Schemes showing a left lateral view of the reedfish brain with the location of the habenulae in red and dotted lines indicating section planes at anterior, medial and posterior levels (a1), and a section at a medial organ level with tract zones in gray (a2). **b-j** Transverse sections of reedfish juvenile habenulae after ISH with probes for *EcSox1* (b,e1-3), *EcNtng2* (c), *EcPcdh17* (d), *EcKctd12a* (f1-3), *EcKiss1* (g1-3), *EcProx1* (h), *EcKctd8* (i) and *EcKctd12b* (j), dorsal to the top. Sections (b-d) were obtained from the same specimen, same for sections (e-g) and sections (h-j). (e1,f1,g1), (e2,f2,g2) and (e3,f3,g3) respectively show sections at anterior, medial and posterior organ levels. Black and white arrowheads point to ventral territories respectively co-expressing *EcSox1/EcNtng2/EcPcdh17* on the left and expressing *EcKiss1* on the right. Thin arrows indicate their boundary with *EcKctd8/12a*-positive dorsal habenula territories. *EcProx1* expression is restricted to the right but spans both dorsal and ventral subdomains, except for a minor dorsal territory, labeled by an asterisk (h). Abbreviations: L, left; R, right. Scale bar=100µm.

**Supplementary Figure 9. Subdomain organization of habenulae in the spotted gar *Lepisosteus oculatus*. a** Schemes showing a left lateral view of the spotted gar brain with the location of the habenulae in red and dotted lines indicating section planes at anterior, medial and posterior levels (a1), and a transverse section at a medial organ levelwith tract zones in gray (a2) . **b-c** Adjacent horizontal sections of habenulae from the same spotted gar juvenile after ISH with probes for *LoPcdh17* (b) and *LoProx1* (c), anterior to the top. **d-k** Transverse sections after ISH with probes for *LoKctd8* (d1-3), *LoKctd12a* (e1-3), *LoKctd12b* (f1-3), *LoKiss1* (g1-3), *LoProx1* (h1-3), *LoSox1* (i1-2), *LoNtng2* (j), and *LoPcdh17* (k1-2), dorsal to the top. Sections (d,e,g) were obtained from the same specimen, same for sections (j,k). (b,c), (d1,e1,f1,g1,h1,i1,i2,j,k1,k2), (d2,e2,f2,g2,h2) and (d3,e3,f3,g3,h3) respectively show sections at ventral, anterior, medial and posterior organ levels. Black and white arrowheads point to bilateral ventral territories respectively co-expressing *LoSox1/LoNtng2/LoPcdh17* and *LoKiss1/LoProx1*. Thin arrows indicate their boundary with *LoKctd8/12b*-positive dorsal habenula territories. *LoNtng2* expression includes a dorsal territory, labeled by an asterisk in (j), in addition to its ventral one. Dotted lines in (d,e) delimit two subdomains, respectively positive or negative for *Kctd12a*, within the dorsal habenulae. Abbreviations: Mes, mesencephalon; Tel, telencephalon; L, left; R, right. Scale bar=100µm.

**Supplementary Figure 10. Habenular subdomain organization in the lungfish *Protopterus annectens*. a** Schemes showing a left lateral view of the lungfish brain with the location of the habenulae in red and dotted lines indicating section planes at anterior, medial and posterior levels (a1), and a section at a medial organ level with tract zones in gray (a2). **b-g** Transverse sections of habenulae from lungfish juveniles, after ISH with probes for *PaPcdh17* (b), *PaNtng2* (c), *PaSox1* (d), *PaKctd8* (e), *PaKctd12b* (f), and *PaProx1* (g), dorsal to the top. (c1,d1,e1,f1,g1), (c2,d2,e2,f2,g2) and (c3,d3,e3,f3,g3) respectively show sections at anterior, medial and posterior organ levels. Black and white arrowheads respectively point to left-restricted *PaSox1* and right-restricted *PaProx1* lateral territories. Thin arrows indicate their boundaries with the *PaKctd8*-positive medial territory. White dotted lines in (d,e) delimit two subdomains, respectively positive or negative for *Kctd12b*, within the *PaKctd8*-positive medial habenula. Abbreviations: hc, habenular commissure; Mes, mesencephalon; Tel, telencephalon; L, left; R, right. Scale bar=100µm.

**Supplementary Figure 11. Subdomain organization of habenulae in the Western clawed frog *Xenopus tropicalis*. a** Schemes showing a left lateral view of the frog brain with the location of habenulae in red and dotted lines indicating section planes at anterior, medial and posterior levels (a1), and a section at a medial organ level with tract zones in gray (a2). **b-j** Transverse sections of frog juvenile habenulae after ISH with probes for *XtSox1* (b,e), *XtNtng2* (c), *XtPcdh17* (d), *XtKctd8* (f,h), *XtKctd12a* (i), *XtKctd12b* (j), and *XtProx1* (g), dorsal to the top. Sections (b,c,d) were obtained from the same specimen, same for sections (e,f,g) and sections (h,i,j). (e1,f1,g1,h1,i1,j1), (b1,c1,d1,e2,f2,g2) and (b2,c2,d2,e3,f3,g3,h2,i2,j2) respectively show sections at anterior, medial and posterior organ levels. Black and white arrowheads respectively point to a bilateral lateral habenula territory co-expressing *XtSox1/XtNtng2* (but not *XtPcdh17*), and an adjacent one expressing *XtProx1*, located more ventrally and excluded from anterior-most levels. Thin arrows indicate the medial/dorsal boundary of the *XtSox1/XtNtng2* territory with medial *XtKctd8/12b*-positive habenula territories. In addition to its habenular territory, *XtSox1* expression expands into a

*XtPcdh17/XtProx1*-positive thalamic territory labeled by an asterisk at medial to posterior levels (b2,d1,d2,e3,g2,g3). Dotted lines in (b,c,e,g) delineate the boundary between the *XtSox1/XtNtng2* and the *XtProx1* territories. Dashed lines in (h1,j1) delimit two anterior subdomains, respectively expressing *XtKctd8* and *XtKctd12b*, within the medial habenulae. Abbreviations: Mes, mesencephalon; Tel, telencephalon; L, left; R, right. Scale bar=50μm.

**Supplementary Figure 12. Heterogeneity of β-catenin distribution in the catshark medial habenula at stage 31. a-f** Confocal images of transverse sections of developing catshark habenulae (stage 31) after HCR-FISH with signals for *ScSox1*, *ScKctd12b* and *ScProx1* in respectively yellow, magenta, and cyan (a,c,e), and after IHC using an antibody directed against β-catenin (red) (b,d,f). DAPI-stained nuclei are shown in blue. All sections were obtained from the same embryo. (a,b), (c,d) and (e,f) each show adjacent sections, at anterior, medial and posterior levels respectively. (b1-2), (d1-8) and (f1-2) show magnifications of the territories boxed in (b), (d) and (f), with β-catenin signals in (b1,d1,d3,d5,d7,f1) and DAPI signals in (b2,d2,d4,d6,d8,f2). White and opened arrowheads respectively point to Left- and Right-LHb, an asterisk in (e) marks a minor *ScProx1* territory on the left. Thin white arrows in (f1-2) point to β-catenin labeled nuclei. A dotted line in (d) delimits a medial territory devoid of cytoplasmic β-catenin signal (d1,d2), similar to the anterior left habenula (b). Based on their location, these territories correspond to the *ScPde1a*-expressing MHb subdomain. A cytoplasmic (but not nuclear) β-catenin signal is observed in all other MHb territories (b1-2;d3-8). Abbreviations: hc, habenular commissure; L, left; R, right. Scale bars=100μm.

**Supplementary Figure 13. Phenotypes observed along the antero-posterior axis of catshark stage 31 habenulae following IWR-1 treatment. a-l** Transverse sections of catshark stage 31 habenulae from control (a-d) and IWR-1-treated (e-h) embryos, after IHC with an antibody directed against β-catenin with β-catenin in red and DAPI-stained nuclei in blue (a,e), and after ISH with probes for *ScSox1* (b,f), *ScProx1* (c,g), *ScRorα* (d,h). Section levels along the antero-posterior axis are indicated at the bottom of each panel (a1-g1, anterior; a2-g2,d1,d3,h1, medial: a3-g3,d2,h2, posterior), dorsal is to the top. (b,c) were obtained from the same embryo, same for (f,g). (b1,c1) are adjacent sections, same for (b2,c2), (b3,c3), (f1,g1), (f2,g2) and (f3,g3). (a1',a1"), (a2',a2"), (a3',a3"), (e1',e1"), (e2',e2"), and (e3',e3") show magnifications of the territories boxed in (a1), (a2), (a3), (e1), (e2), and (e3) respectively, with β-catenin signals in (a1',a2',a3',e1',e2',e3') and DAPI signals in (a1",a2",a3",e1"e2",e3"). Arrowheads in (e-g) show a loss of nuclear β-catenin accumulation in the lateral right habenula (e) and an expansion of *ScSox1* expression to this territory, concomitant with a loss of *ScProx1* expression. Asterisks in (d1,d2) indicate lateral right-restricted territory lost in IWR-1 treated embryo (h1,h2). Abbreviations: L, left; R, right. Scale bar=100µm.

**Supplementary Figure 14. IWR-1 treatment at stage 29 has no effect on *ScPde1a* and *ScEnpp2* expression in developing catshark habenulae. a-d** Transverse sections of catshark stage 31 habenulae from control (a-b) and IWR-1 treated (c-d) embryos, after ISH with probes for *ScPde1a* (a,c) and *ScEnpp2* (b,d). Sections (a1 to a4), (b1 to b4), (c1 to c4), and (d1 to d4) progress along the antero-posterior axis of the habenulae from anterior to posterior levels, dorsal is to the top. (a,b) are adjacent sections from the same embryo, same for (c,d). (a1,b1) are adjacent sections, same for (a2,b2), (a3,b3), (a4,b4), (c1,d1), (c2,d2), (c3,d3) and (c4,d4). Dotted lines delimit internal and external components of the medial habenula, a thin arrow points to the boundary between the complementary *ScPde1a* and *ScEnpp2* territories in the left external medial habenula. Black and white arrowheads show the location of left and right lateral habenulae respectively. The number "n" refers to the number of embryos taken into account and showing the same asymmetry pattern for these makers. Abbreviations: L, left; R, right. Scale bar=100µm.

**Supplementary Figure 15. Phenotypes observed along the antero-posterior axis of catshark stage 31 habenulae following double SB-505124 and IWR-1 treatment. a-h** Transverse sections of catshark stage 31 habenulae from SB-505124 (a-d) and double SB-505124/IWR-1-treated (e-h) embryos, after IHC with an antibody directed against β-catenin with β-catenin in magenta and DAPI-stained nuclei in blue (a,e), HCR-FISH with territories of *ScSox1* and *ScProx1* in yellow and cyan, respectively (b,f), and ISH with probes for *ScSox1* (c,g) and *ScProx1* (d,h). Section levels along the antero-posterior axis are indicated at the bottom of each panel (a1-h1, anterior; c2,d2,g2,h2, medial: a2,b2,c3,d3,e2,f2,g3,h3, posterior), dorsal to the top. (a,b) were obtained from the same embryo, same for (c,d), (e,f) and (g,h). (c1,d1) are adjacent sections, same for (c2,d2), (c3,d3), (g1,h1), (g2,h2) and (g3,h3). A right isomerism is observed at all axial levels in SB-505124 treated embryos (a-d). Arrowheads in (e-h) point to bilateral territories where nuclear β-catenin accumulation (e) and *ScProx1* expression (f,h) are lost, and where *ScSox1* expression is rescued in the lateral habenulae of double SB-505124/IWR-1-treated embryos. These territories are never observed in embryos only treated with SB-505124 (a-d). Scale bar=100µm.

**Supplementary Figure 16. Spatial and temporal regulation of progenitor cycle exits in the developing catshark habenulae. a-f** Horizontal sections of catshark stage 31 habenulae following exposure of embryos to BrdU pulses at stage 28 (a,b), 28+ (c,d), and 29 (e,f), anterior to the top. (a,c,e) show confocal images following IHC using an antibody directed against BrdU (green; DAPI-stained nuclei in gray). (b,d,f) respectively show confocal images after double ISH with probes for *ScKctd12b* (magenta) and *ScProx1* (cyan), with DAPI-stained nuclei in gray. (a1,b1), (a2,b2), (a3,b3) each show adjacent sections of the same embryo at anterior, medial and posterior levels respectively, same for (c1,d1), (c2,d2), (c3,d3), and (e1,f1), (e2,f2), (e3,f3). White dotted lines in (b,d,f) delimit the border between medial (MHb) and lateral (LHb) habenular territories, as inferred from *ScKctd12b* expression. The approximate location of this border is also shown on adjacent BrdU labeled sections (a,c,e). Thin arrows point to BrdU negative territories in the differentiating habenulae. Abbreviations: hc, habenular commissure; L, left; R, right. Scale bar=100µm.

**Supplementary Figure 17. Asymmetric β-catenin signals persist in the habenulae of catshark juveniles. a-c** Confocal images of transverse sections of catshark juvenile habenulae after IHC using an antibody directed against β-catenin, dorsal to the top. Sections in (a), (b), and (c) are shown at anterior, medial, and posterior levels respectively. DAPI-stained nuclei are shown in blue. All sections were obtained from the same embryo. (a1-8), (b1-4) and (c1-6) show magnifications of the territories boxed in (a), (b) and (c), with DAPI signals in (a1,a3,a5,a7,b1,b3,c1,c3,c5) and DAPI/β-catenin signals in (a2,a4,a6,a8,b2,b4,c2,c4,c6). White arrowheads and black arrowheads point to Left- and Right-LHb respectively. Abbreviations: L, left; R, right. Scale bar=100µm.

**Supplementary Table 1. Transcriptomic analysis of habenular asymmetries in the catshark stage 31 habenulae. Sheet 1** List of differentially expressed genes between the left and the right habenula sides. Differentially expressed gene models are listed in column A, with left- and right-enriched ones shaded in yellow and blue respectively. Curated annotation and automated annotation against Swissprot are indicated in columns B and C. P-values, q-values and fold changes (FC) are given in columns D, E and G. Genes previously identified as left-enriched in the catshark developing habenulae (Lagadec et al, 2015. *Nature com.* 6, 6686) such as *ScPitx2* and *ScKctd12b,* are present in this list. **Sheet 2** Functional annotation of left- and right enriched genes. GO (gene ontology) terms for biological processes over-represented in the lists of left- and right-enriched genes are represented in columns (B,C) and (G,H) respectively, with the corresponding q-values in A and F, and the identity of genes taken into account in D and I. Terms are grouped together by level (2 to 5). Those retrieved both for left- and right enriched genes are shown in red characters.

*See accompanying file : Suppl Table1.xls.*

| | Catshark | Mouse | Zebrafish |
|---|---|---|---|
| *Sox1* | Left-LHb | - | Hb11; Ad_Hb11 (*sox1a/b*): *Pandey et al. 2018* |
| *Pcdh17* | | LHb: *Allen Brain Atlas* | - |
| *Lhpfl5* | | - | - |
| *Ntng2* | | LHb, MHb: *Allen Brain Atlas* | - |
| *Ptprm* | | - | - |
| *Spon1* | External-MHb anterior left | lateral MHb: *Allen Brain Atlas* | Hb07 (*spon1a*): *Pandey et al. 2018* |
| *Stk32c* | | - | - |
| *Trhde* | | lateral MHb: *Allen Brain Atlas* | - |
| *Trhr2* | | MHb: *Heuer et al. 2000* | - |
| *Pde1a* | | MHb: *Allen Brain Atlas* | - |
| *Kctd12a* | | MHb: *Metz et al. 2011* | Ad_Hb04/06 (*kctd12.1, lov*), Ad_Hb02B (*kctd12.2, ron*): *Pandey et al. 2018* |
| *Kctd12b* | MHb | MHb: *Metz et al. 2011* | - |
| *Kctd8* | | MHb*: Allen Brain Atlas, Metz et al. 2011* | - |
| *Nrp2* | | Hb: *Allen Brain Atlas* | - |
| *Stac1* | | MHb: *Allen Brain Atlas* | - |
| *Enpp2* | External MHb, right + posterior left | - | - |
| *Eya4* | | - | - |
| *Ak5* | Right-LHb | - | Hb15; Ad_VHb01 (*ak5*): *Pandey et al. 2018* |
| *Kiss1* | | - | Hb15; Ad_VHb01 (*kiss1*): *Pandey et al. 2018* |
| *Prox1* | | PVT: *Allen Brain Atlas* | Hb01 (*prox1a*): *Pandey et al. 2018* |
| *Rerg* | | PVT: *Allen Brain Atlas* | Ad_VHb02/03/04 : *Pandey et al. 2018* |
| *Stxbp6* | | Thalamic nuclei adjacent to HB, including PVT: *Allen Brain Atlas* | - |
| *Prkcq* | | MHb+dispersed LHb cells : *Allen Brain Atlas* | Hb15 (*prkcq*): *Pandey et al. 2018* |
| *Wisp1* | | - | - |
| *Gng14* | | - | Hb15; Ad_VHb01 (si:dkey-117i10.1): *Pandey et al. 2018* |
| *Shh* | | - | - |
| *Trib1* | | - | - |

**Supplementary Table 2. Expression characteristics of mouse and zebrafish orthologs of markers of the main habenular territories identified in the catshark.**

**Supplementary Table 2. Expression characteristics of mouse and zebrafish orthologs of markers of the main habenular territories identified in the catshark.** Markers of the broad territories identified in the catshark habenulae are listed in the first column, with their expression site in the second column. Expression territories of their orthologs in the mouse habenulae or adjacent thalamic territories are shown in the third column, based on references cited or searches in the Allen Brain Atlas. The fourth column shows the zebrafish orthologs, which were identified as signatures of cell clusters in a single-seq RNA-seq characterization of the zebrafish habenulae (Pandey et al. 2018). The identity of cell clusters is indicated and their location in the habenulae is the following: Hb11 and Ad_Hb11, larval and related adult ventral cell clusters; Hb07, left-enriched dorsal cell cluster; Ad-Hb02B/04/06, adult cell clusters related to larval dorsal hebenula cell clusters (right enriched for Ad_Hb02B); Hb15, larval ventral cell cluster; Hb01, larval dorsal right enriched cell cluster; Ad_VHb01/02/03/04, adult ventral cell clusters. References cited: Allen Brain Atlas, Lein et al. 2007. Nature 445:168-176; Heuer et al. 2000. J. Comp. Neurol. 428:319-336; Metz et al. 2011. J. Comp. Neurol. 519:1435-1454; Pandey et al. 2018. Curr Biol. 8:1052-1065.e7.

| Phenotype / Marker | Control embryos | | IWR-1-treated embryos | |
|---|---|---|---|---|
| | Same phenotype as untreated embryos | Presence of zones of Left-LHB identity on the right | Same phenotype as untreated embryos | Presence of zones of Left-LHB identity on the right |
| *ScSox1* | 4/4 | 0/4 | 0/5 | 5/5 |
| *ScNtng2* | 2/2 | 0/2 | 0/2 | 2/2 |
| β–catenin | 4/4 | 0/4 | 0/4 | 4/4 |
| *ScProx1* | 4/4 | 0/4 | 0/5 | 5/5 |
| *ScKiss1* | 1/1 | 0/1 | 0/1 | 1/1 |
| *ScRorα* | 1/1 | 0/1 | 0/1 | 1/1 |
| Total number of embryos | 5/5 | 0/5 | 0/6 | 6/6 |

**Supplementary Table 3. Asymmetry phenotypes in the lateral habenulae of control and IWR-1-treated catshark embryos.** Phenotypes were analyzed at stage 31, following treatments as described in Materials and methods. For each marker listed in the first column, the ratio shown refers to the number of embryos devoid of, or exhibitinging zones of Left-LHB identity on the right, compared to the number of embryos analyzed. For *ScSox1* and *ScProx1*, embryo counts take into account phenotypes observed after detection both by chromogenic ISH and HCR. The last line refers to the total number of embryos analyzed.

| | SB-505124-treated embryos embryos | | SB-505124/IWR-1-treated embryos | |
|---|---|---|---|---|
| Phenotype<br>Marker | Right isomerism | Presence of bilateral zones of Left-LHB identity | Right isomerism | Presence of symmetric zones of Left-LHB identity |
| *ScSox1* | 4/4 | 0/4 | 0/6 | 6/6 |
| *ScNtng2* | 2/2 | 0/2 | 0/2 | 2/2 |
| β–catenin | 2/2 | 0/2 | 0/3 | 3/3 |
| *ScProx1* | 4/4 | 0/4 | 0/6 | 6/6 |
| *ScKiss1* | 2/2 | 0/2 | 0/2 | 2/2 |
| Total number of embryos | 5/5 | 0/5 | 0/7 | 7/7 |

**Supplementary Table 4. Asymmetry phenotypes in the lateral habenulae of SB-505124- and SB-505124/IWR-1-treated catshark embryos.** Phenotypes were analyzed at stage 31, following treatments as described in Materials and methods. For each marker listed in the first column, the ratio shown refers to the number of embryos showing a right isomerism, or exhibitingbilateral zones of Left-LHB identity, compared to the number of embryos analyzed. For *ScSox1* and *ScProx1*, embryo counts take into account phenotypes observed after detection both by chromogenic ISH and HCR. The last line refers to the total number of embryos analyzed.

**Supplementary Table 5. List of genes analyzed by ISH in the catshark *S. canicula* (sheet 1), the elephant shark *C. milii*, reedfish *E. calabaricus*, spotted gar *L. oculatus*, lungfish *P. annectens* and Western clawed frog *X. tropicalis* (sheets 2-6 respectively). Sheet 1** The genes shown were retrieved from the transcriptomic comparison between stage 31 left and right habenulae except for *ScKctd8*, *ScKctd12a* and *ScRorα*, selected by candidate gene approaches as described in Results. Gene names are indicated in column A, the corresponding identification in the reference database in column B, the laterality of the enrichment predicted by the transcriptomic analysis in column C, the rank in the list of differentially expressed genes in column D (ranking on increasing p-values), the corresponding expression characteristics in column E and the sequence of the probe in column F. Colors indicate the expression characteristics of genes exhibiting regionalized ISH in profiles in differentiated habenula territories with the following code: yellow, Left-LHb; light purple, internal MHb; dark purple, external MHB, right and posterior-left; magenta, external MHB, anterior-left; blue, Right-LHb). Genes expressed in combinations of these domains are shaded in grey. **Sheet 2-6** Columns A, B and C respectively contain gene names, their NCBI identifier and probe sequences.

*See accompanying file : Suppl Table5.xls*

a

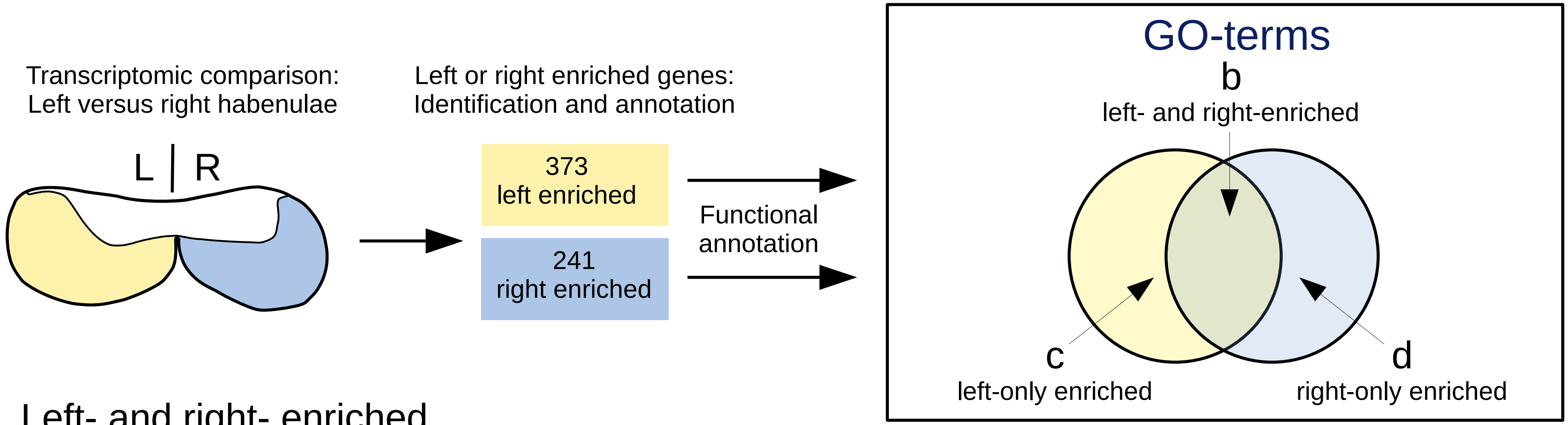


### b Left- and right- enriched

#### CENTRAL NERVOUS SYSTEM

| Term | Left | Right |
|---|---|---|
| neuron differentiation | 7.3E-10 | 3.9E-05 |
| neurogenesis | 2.3E-08 | 1.5E-06 |
| neuron projection guidance | 1.6E-04 | 2.7E-03 |
| neuron projection development | 2.7E-06 | 4.1E-04 |
| synaptic signaling | 6.5E-10 | 3.5E-08 |
| synapse organization | 8.0E-07 | 8.3E-03 |
| regulation of trans-synaptic signaling | 7.1E-09 | 3.0E-05 |
| regulation of membrane potential | 7.4E-04 | 6.1E-03 |

#### ORGANISMAL RESPONSES

| Term | Left | Right |
|---|---|---|
| regulation of locomotion | 3.1E-03 | 6.2E-06 |
| sensory perception of pain | 2.8E-02 | 6.4E-03 |

#### SIGNALING PATHWAYS

| Term | Left | Right |
|---|---|---|
| cell-cell signaling by Wnt | 1.8E-02 | 4.9E-02 |
| glutamate receptor signaling pathway | 1.3E-07 | 3.9E-02 |
| neuropeptide signaling pathway | 2.0E-04 | 4.8E-06 |
| dopamine secretion | 8.9E-04 | 2.7E-02 |

### c Left- only enriched

#### ORGANISMAL RESPONSES

| Term | Left |
|---|---|
| sensory perception of mechanical stimulus | 4.1E-04 |
| feeding behavior | 2.7E-02 |
| sleep | 4.9E-02 |

#### SIGNALING PATHWAYS

| Term | Left |
|---|---|
| adenylate cyclase-modulating GPCR | 2.3E-05 |
| acetylcholine receptor | 1.5E-03 |
| G protein coupled glutamate receptor | 1.5E-03 |
| ionotropic glutamate receptor | 6.8E-03 |

### d Right- only enriched

#### ORGANISMAL RESPONSES

| Term | Right |
|---|---|
| learning or memory | 1.3E-03 |
| response to amphetamine | 2.7E-03 |
| cognition | 4.5E-03 |
| regulation of behavioral fear response | 9.8E-03 |
| behavioral response to nicotine | 9.6E-03 |

#### SIGNALING PATHWAYS

| Term | Right |
|---|---|
| opiod receptor | 2.4E-02 |
| phospholipase C-activating GPCR | 4.8E-02 |

Supplementary Figure 1

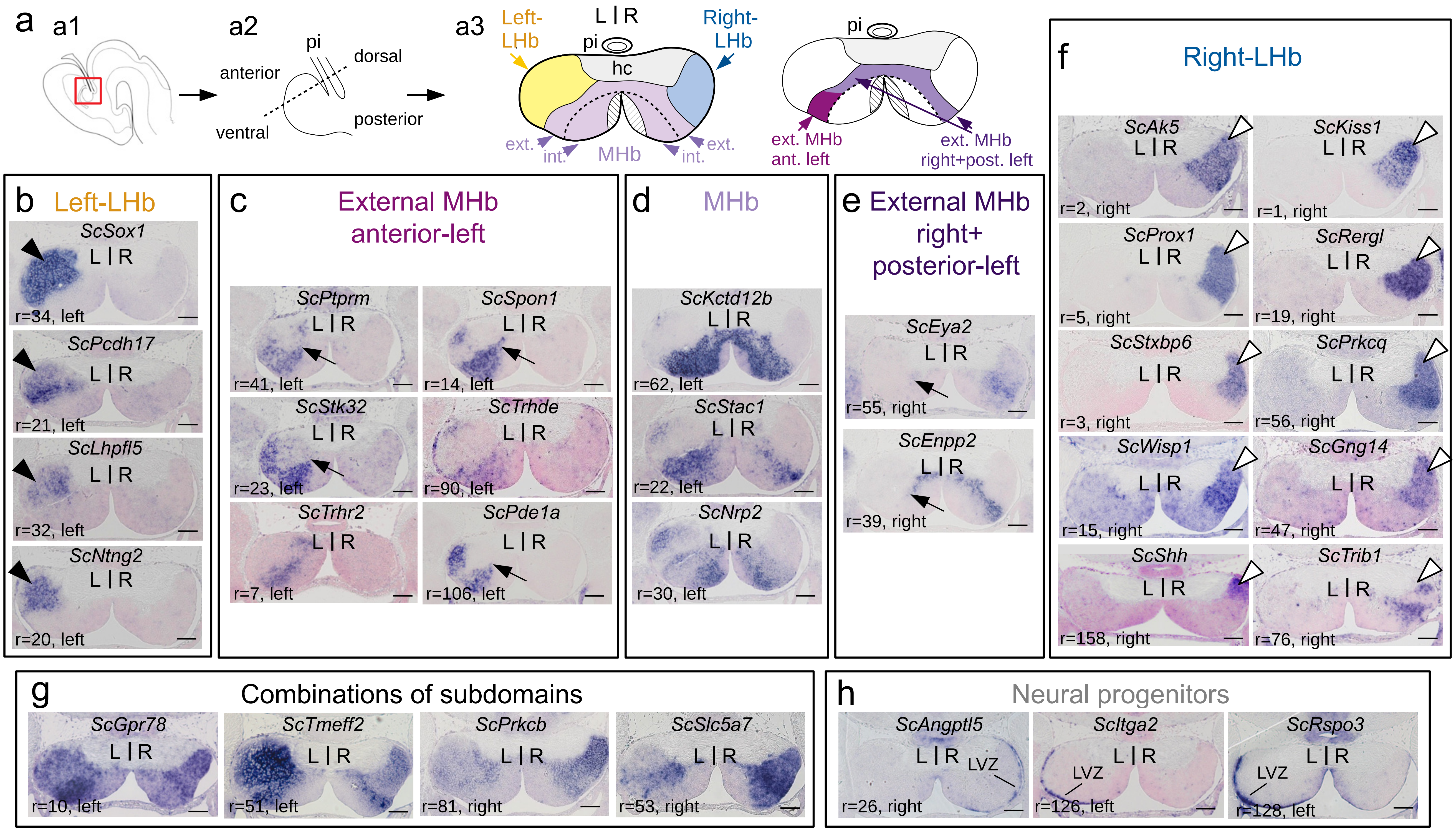


Supplementary Figure 2

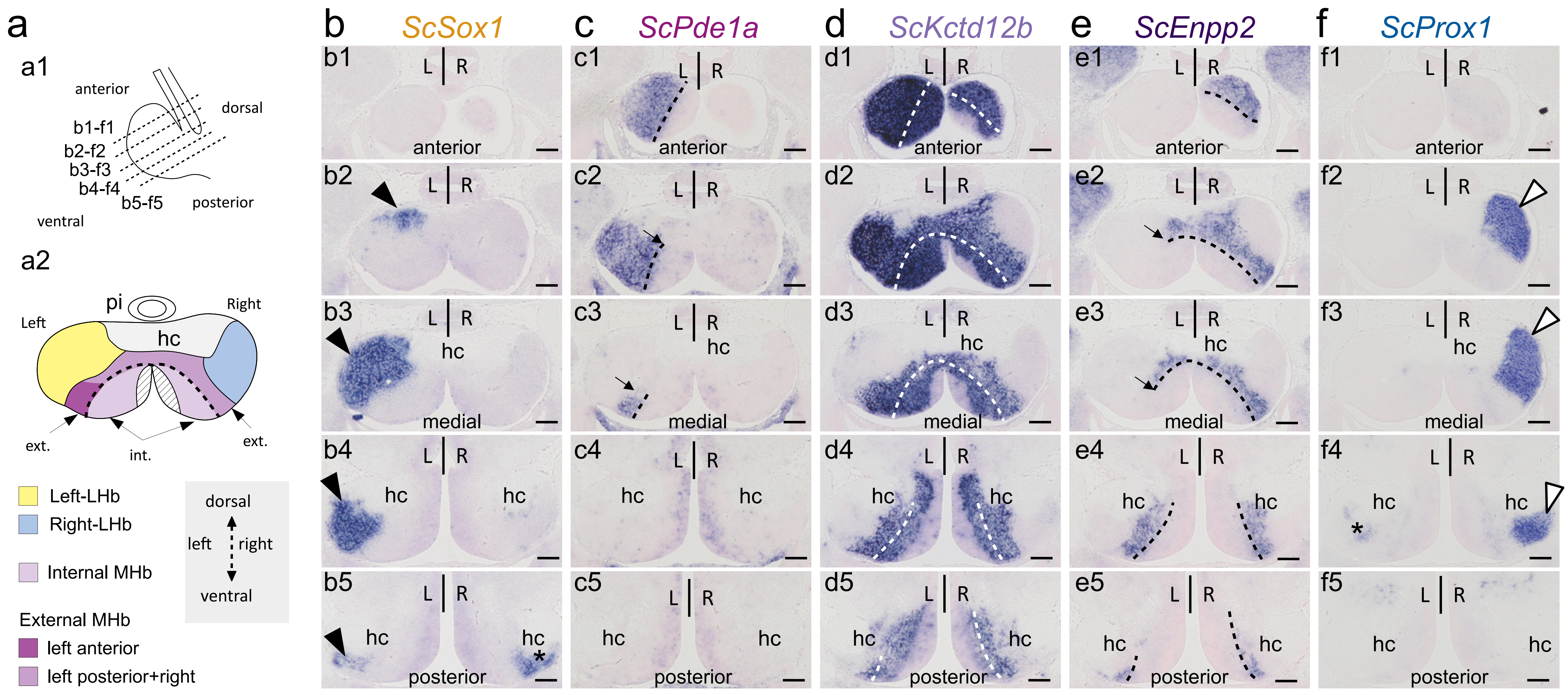


Supplementary Figure 3

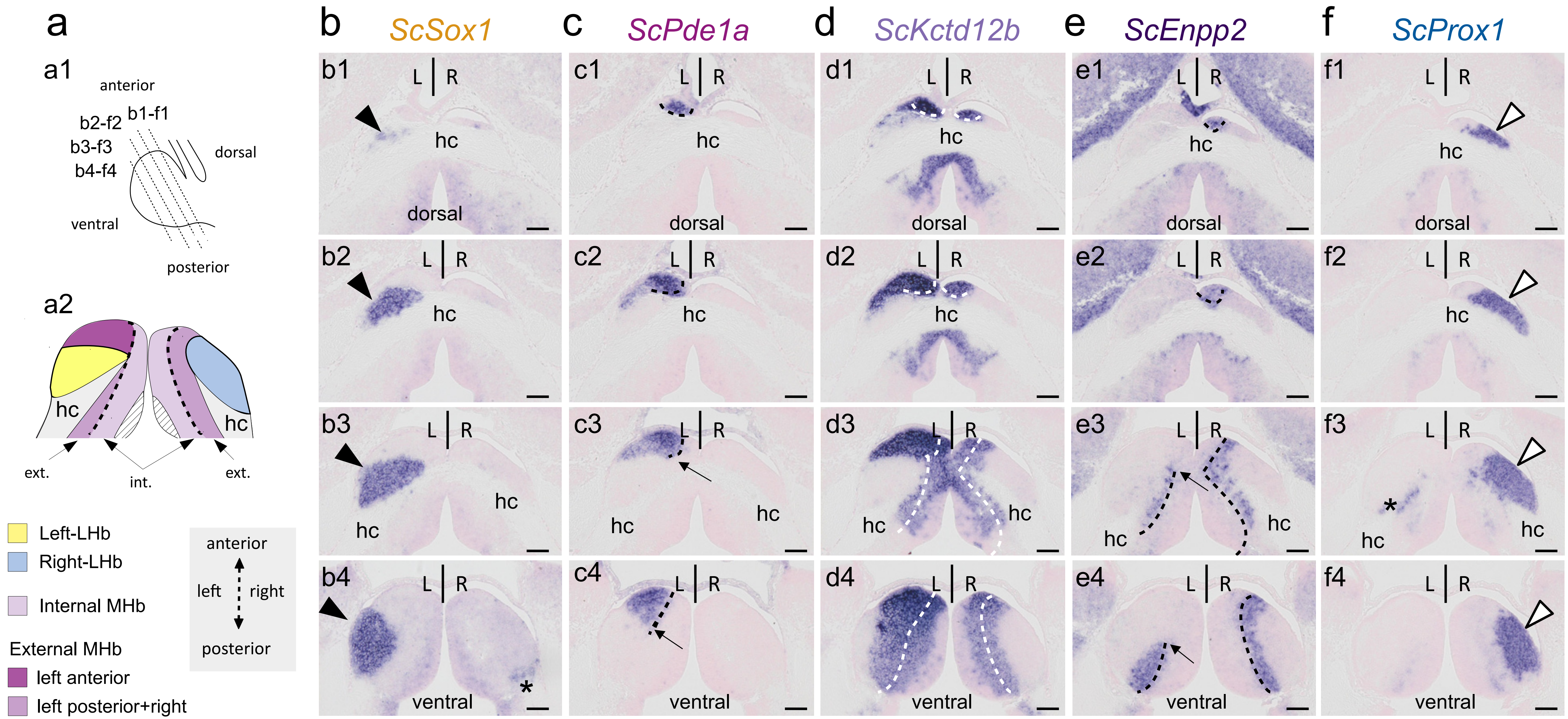


Supplementary Figure 4

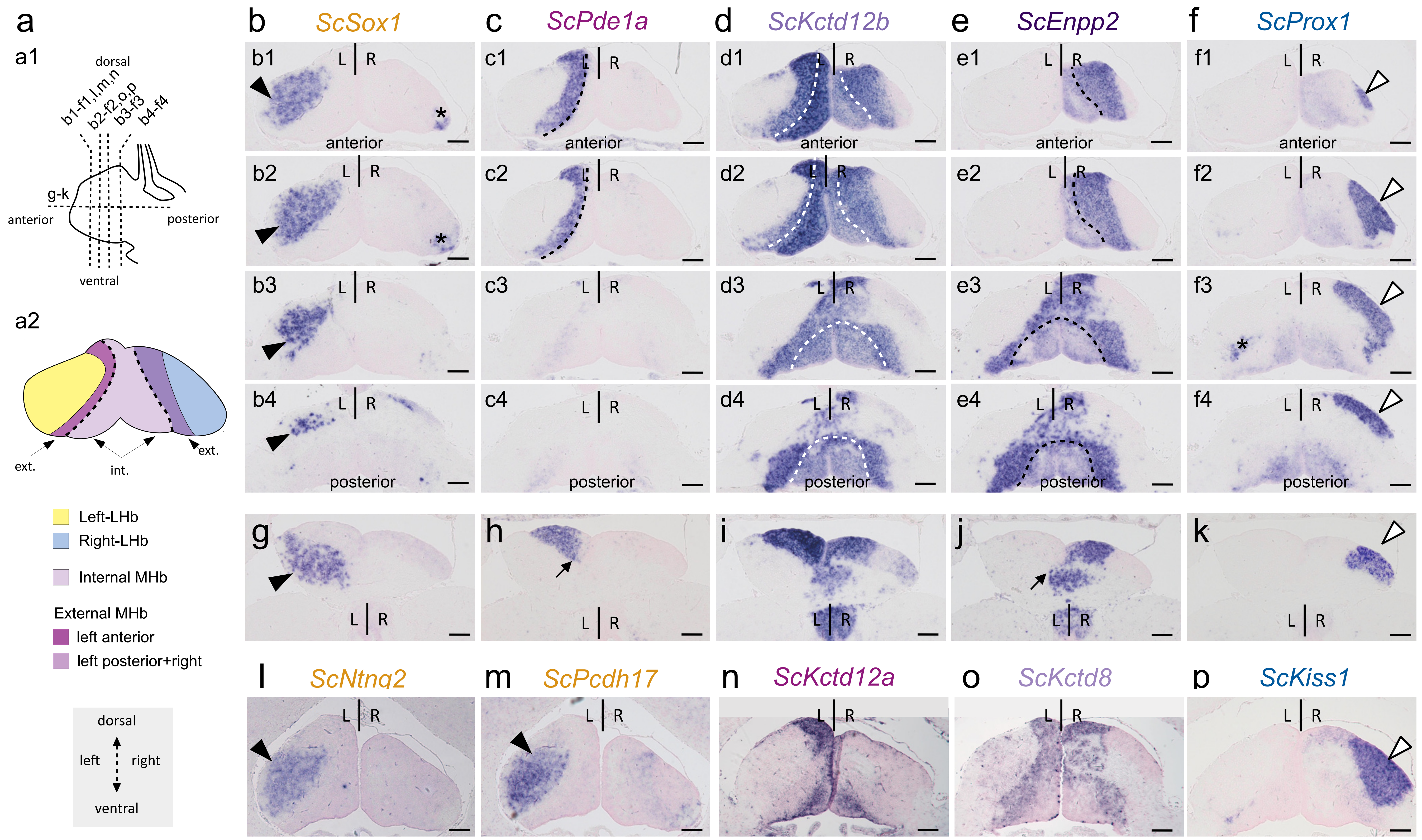


Supplementary Figure 5

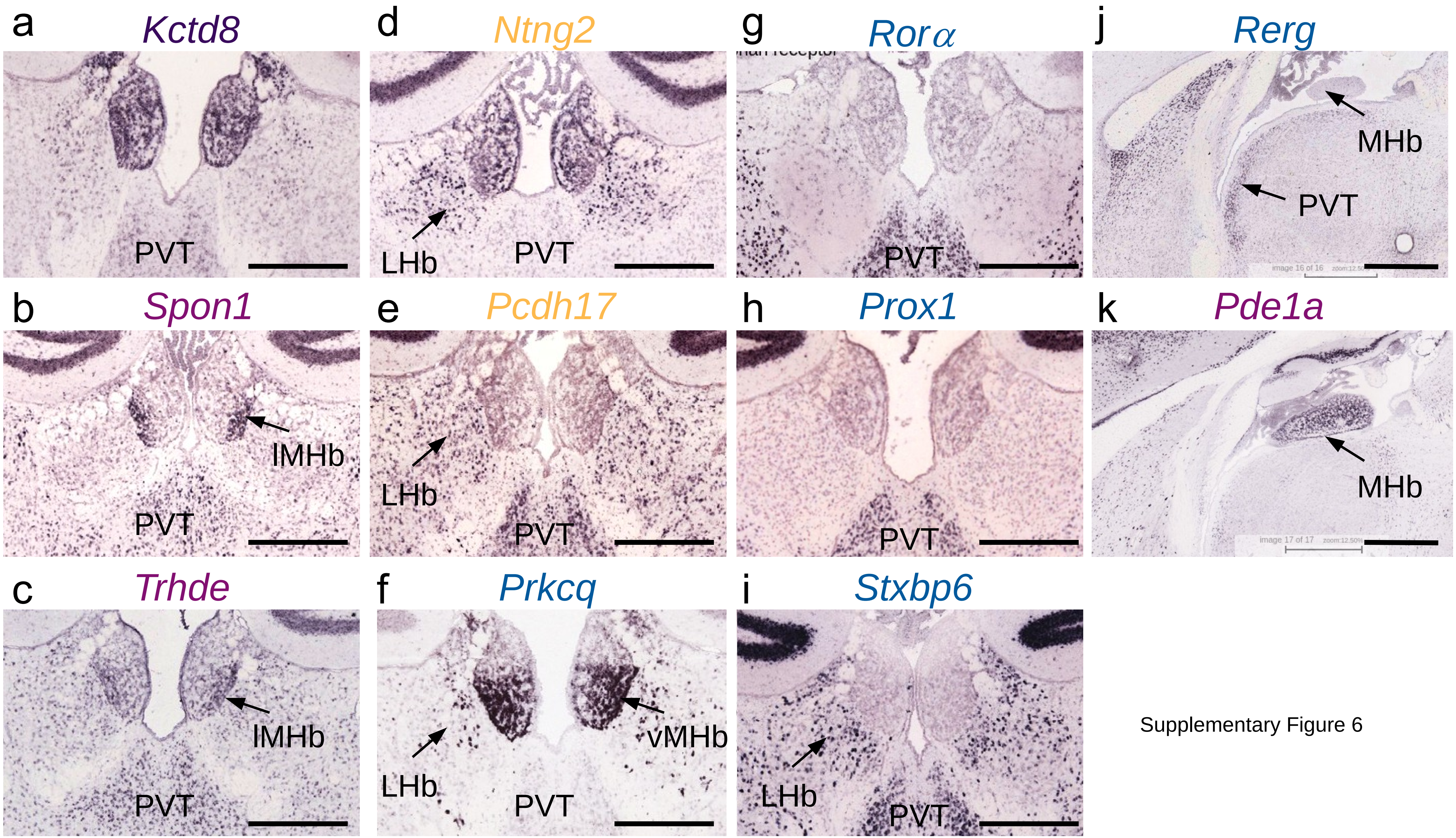


Supplementary Figure 6

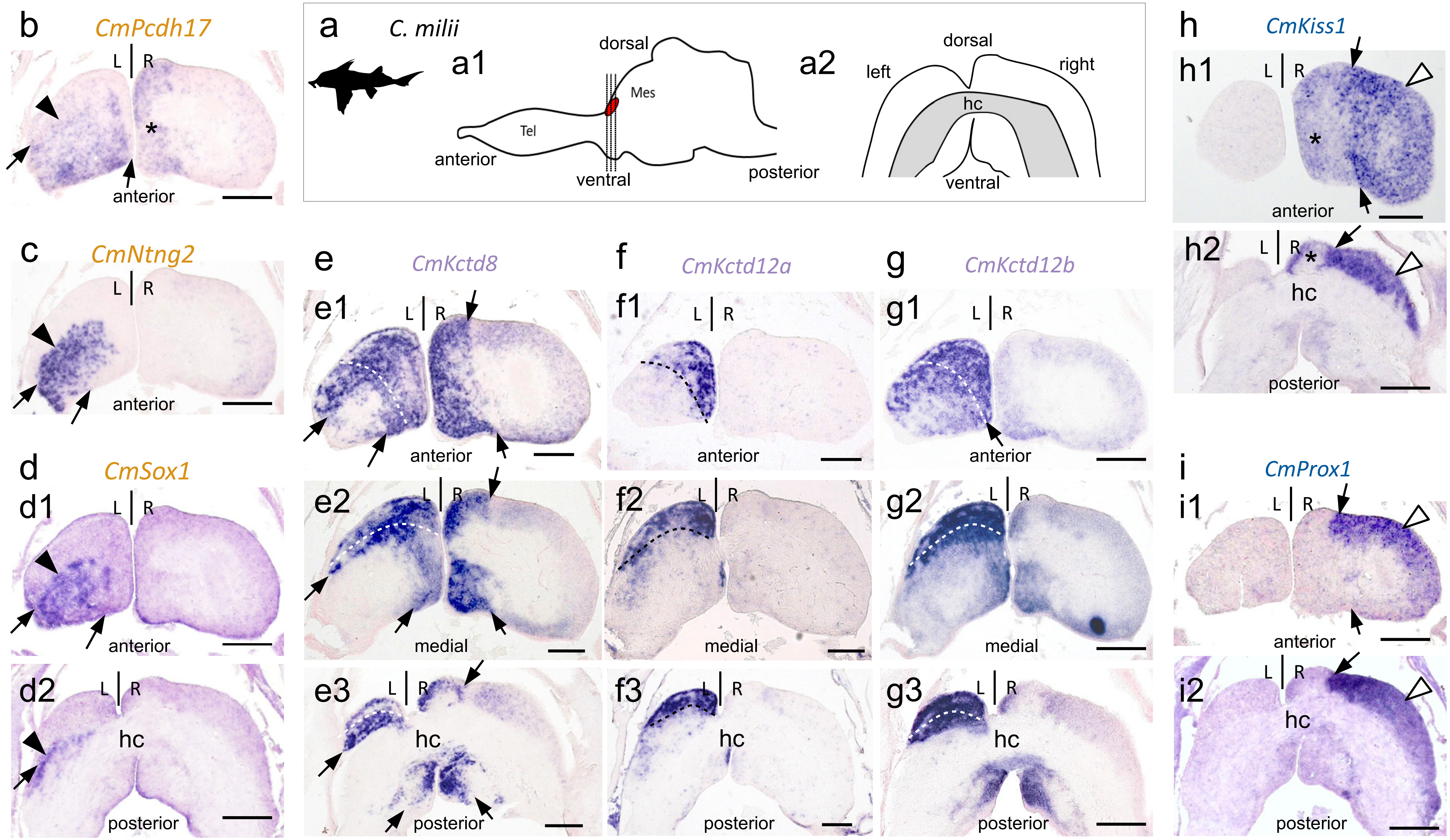


Supplementary Figure 7

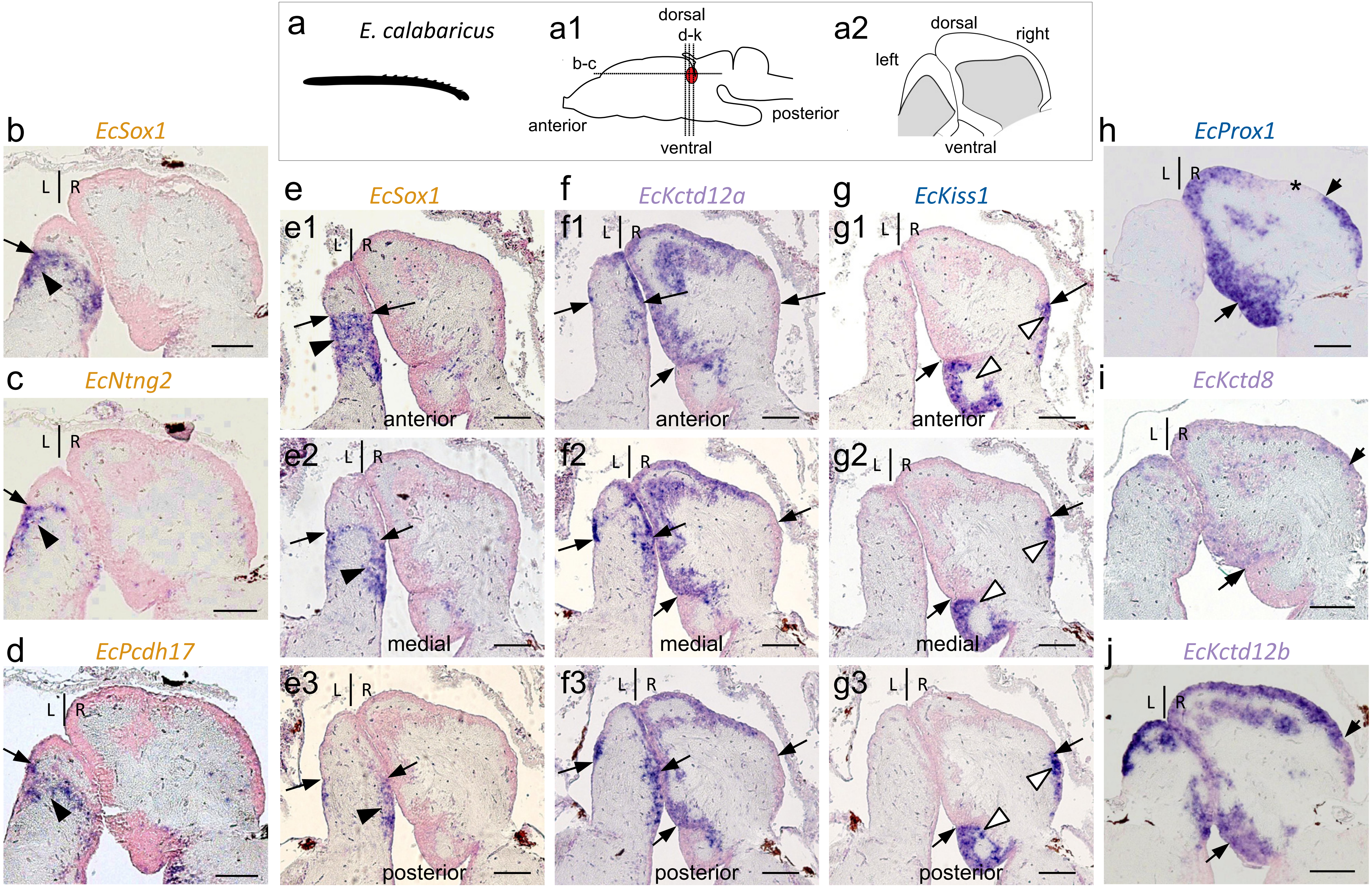


Supplementary Figure 8

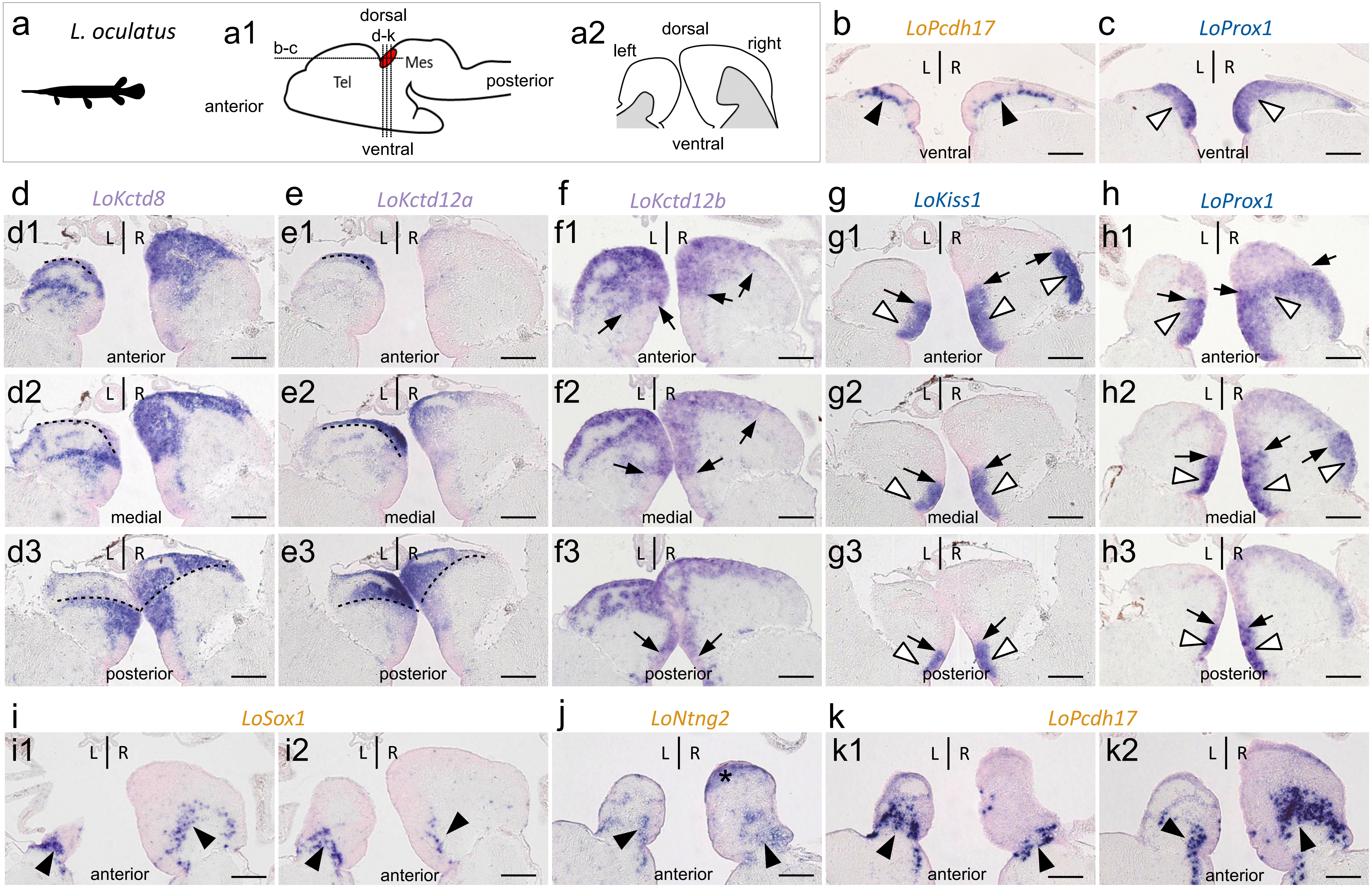


Supplementary Figure 9

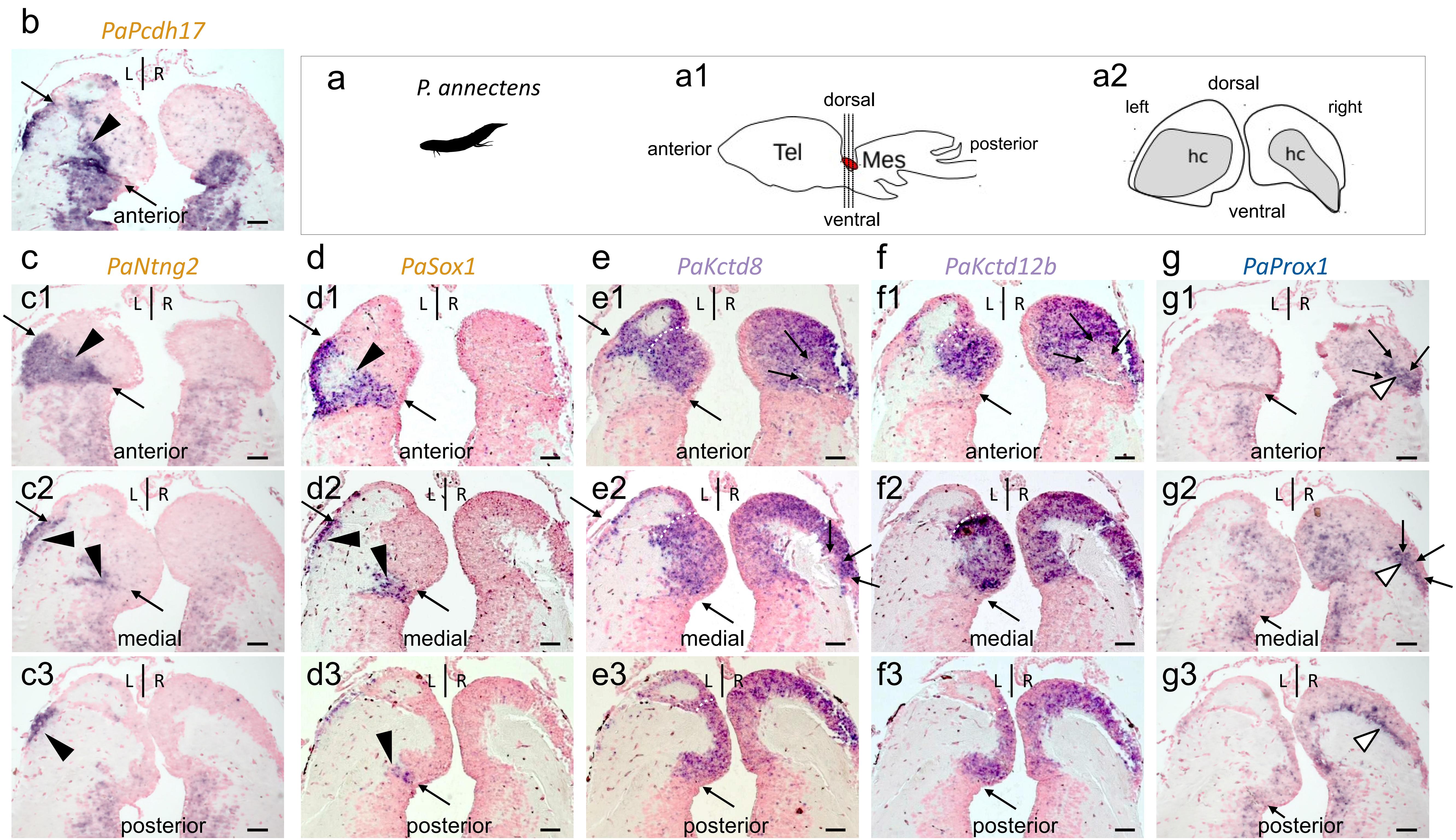


Supplementary Figure 10

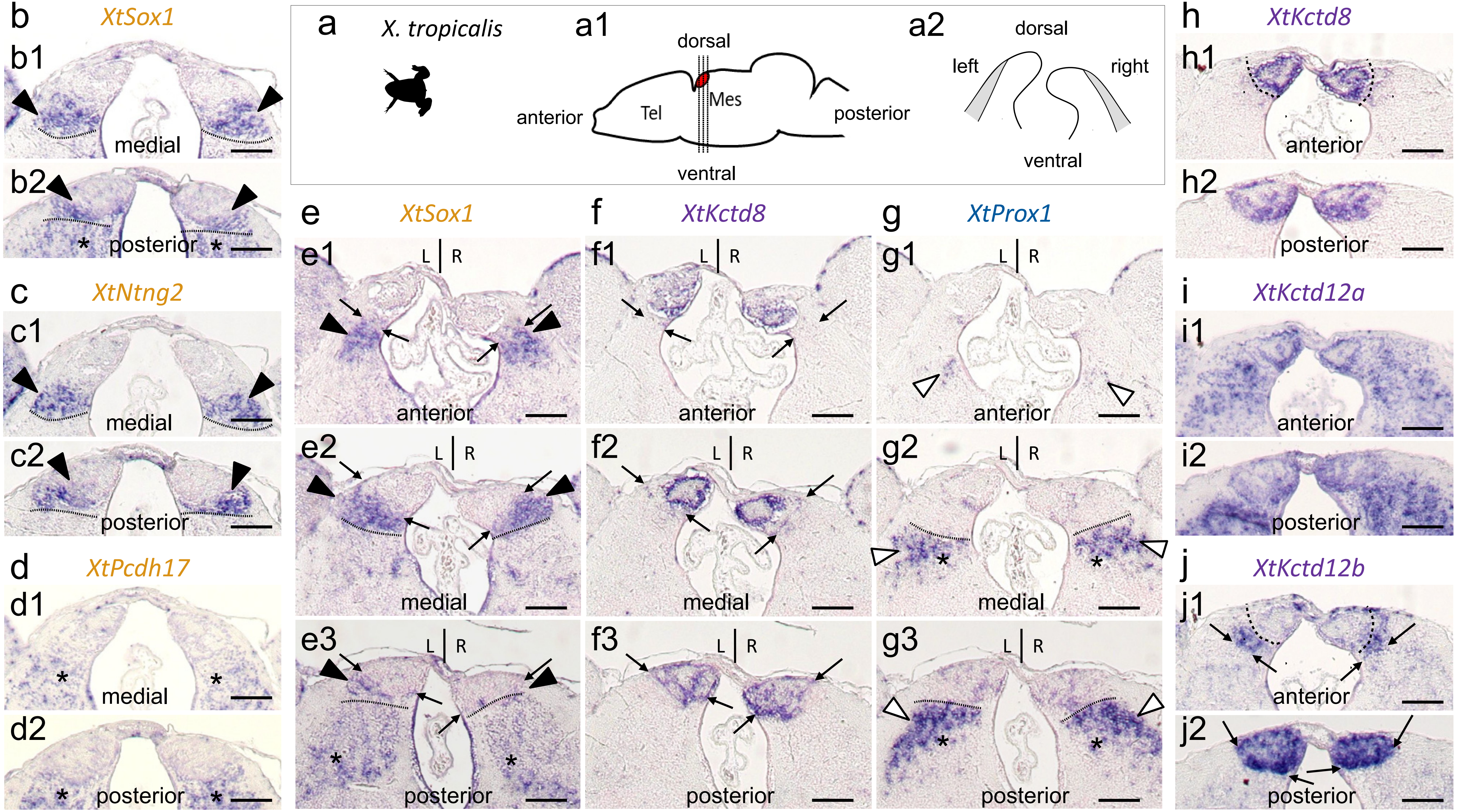


Supplementary Figure 11

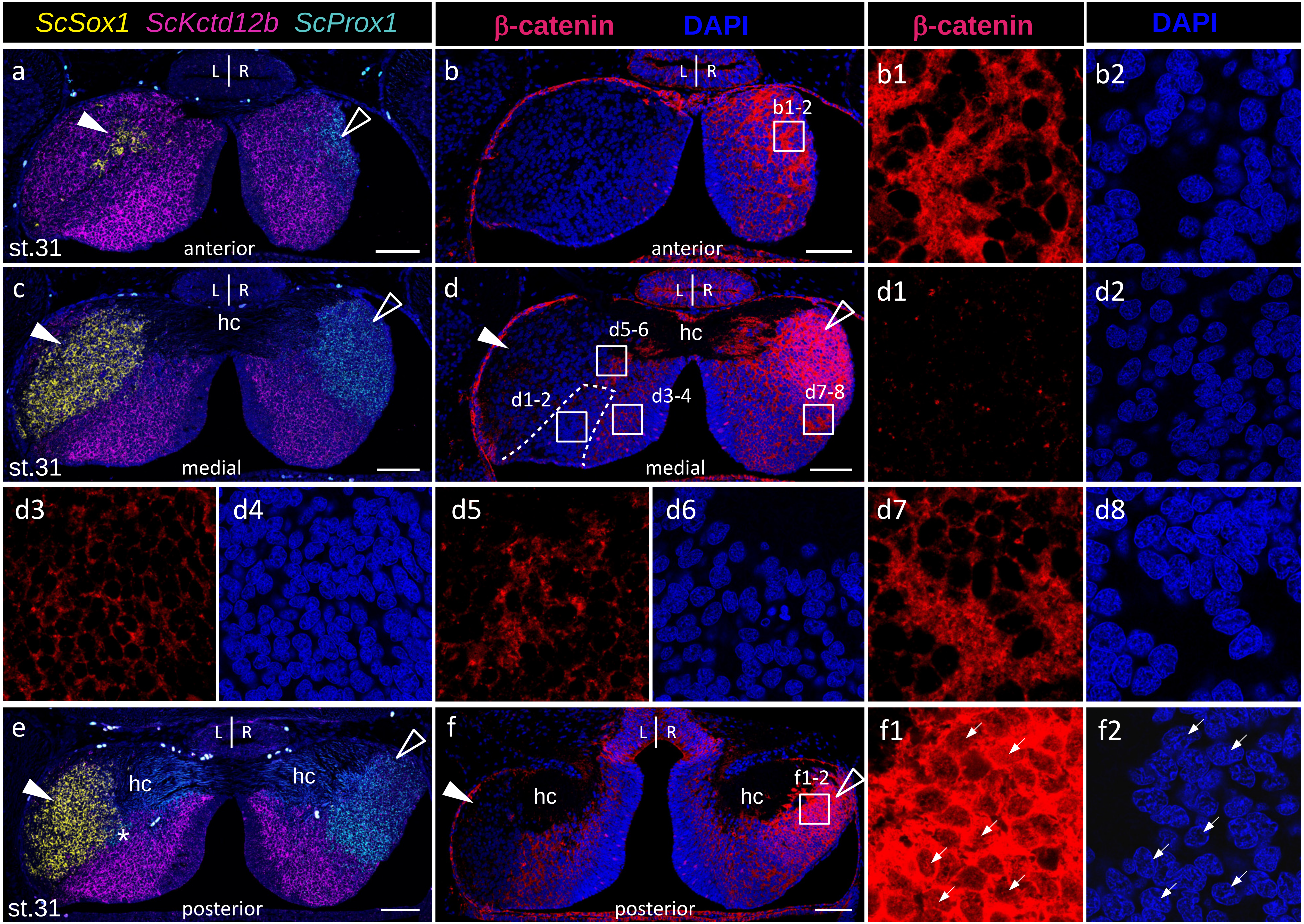


Supplementary Figure 12

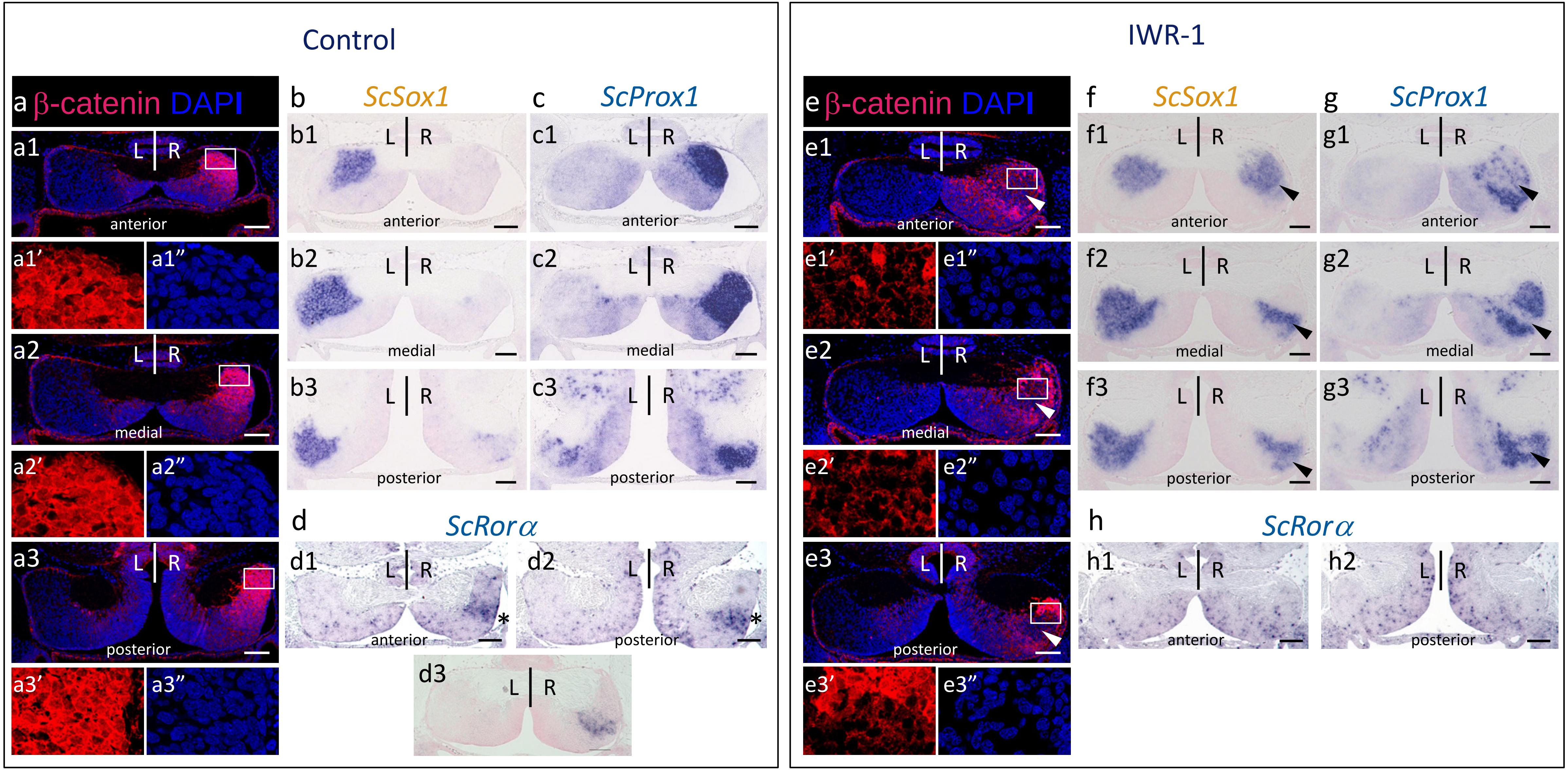


Supplementary Figure 13

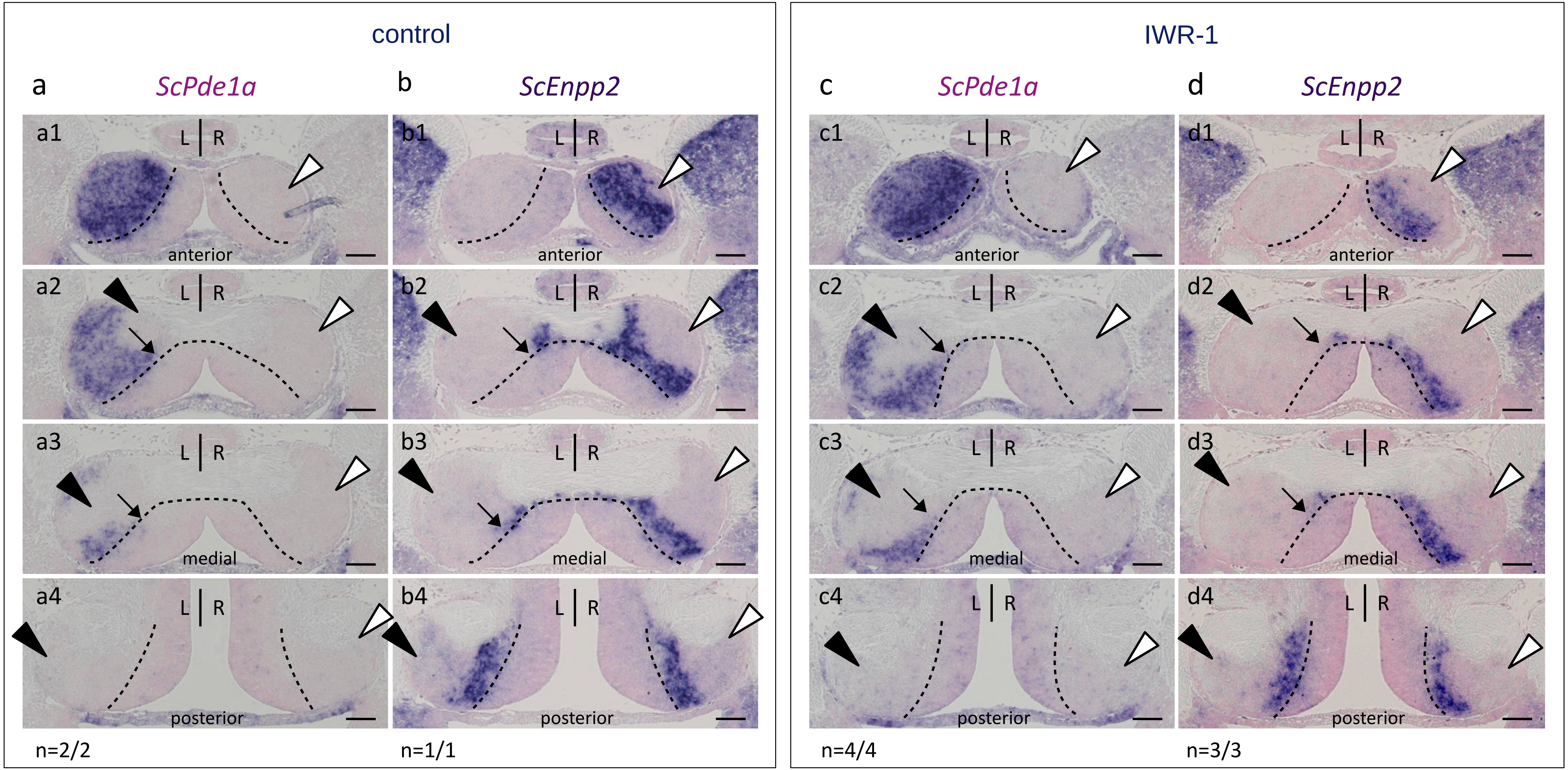


Supplementary Figure 14

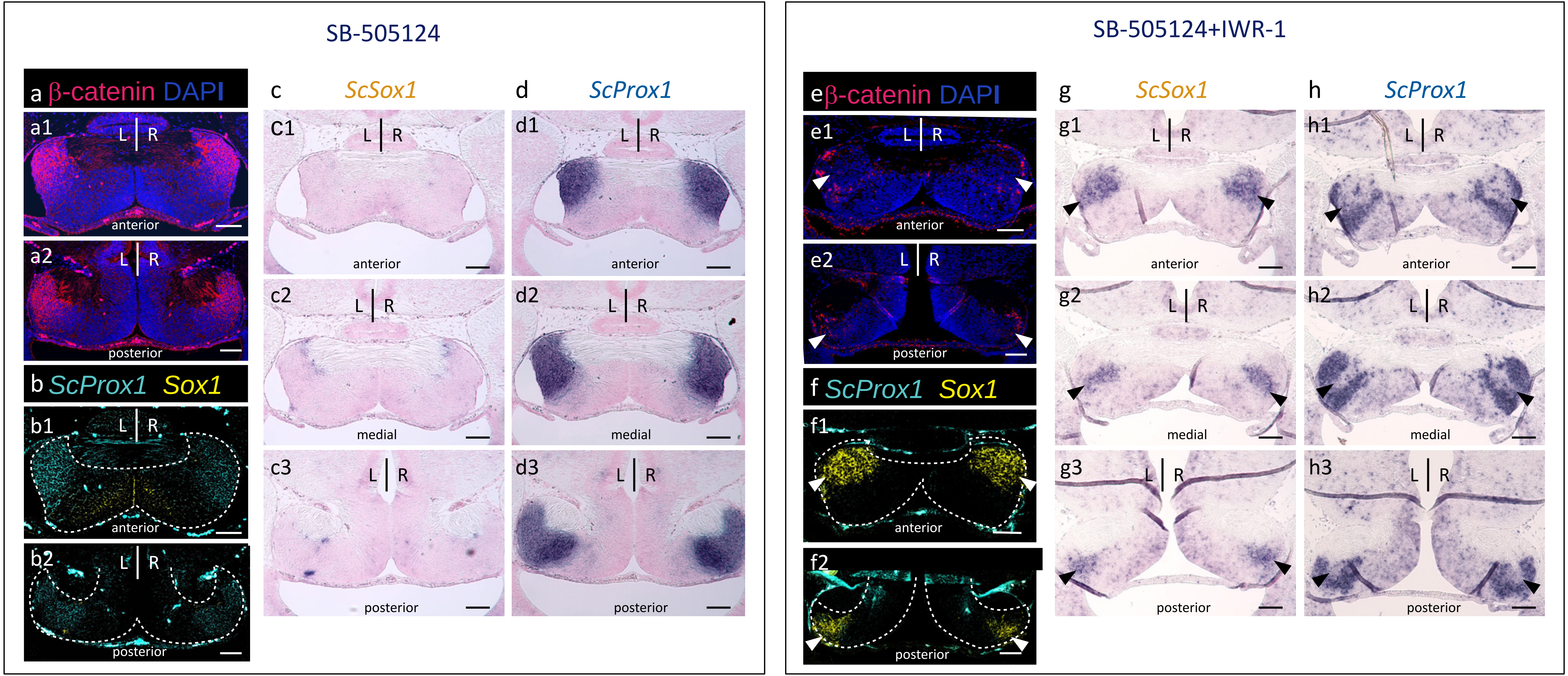


Supplementary Figure 15

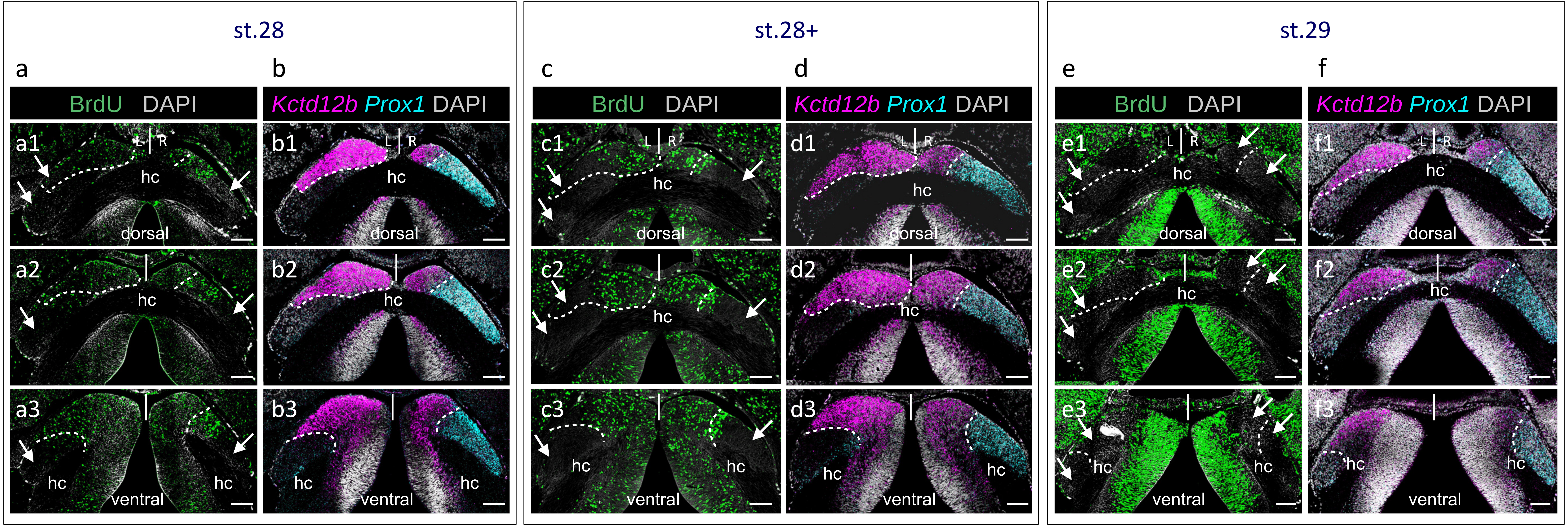


Supplementary Figure 16

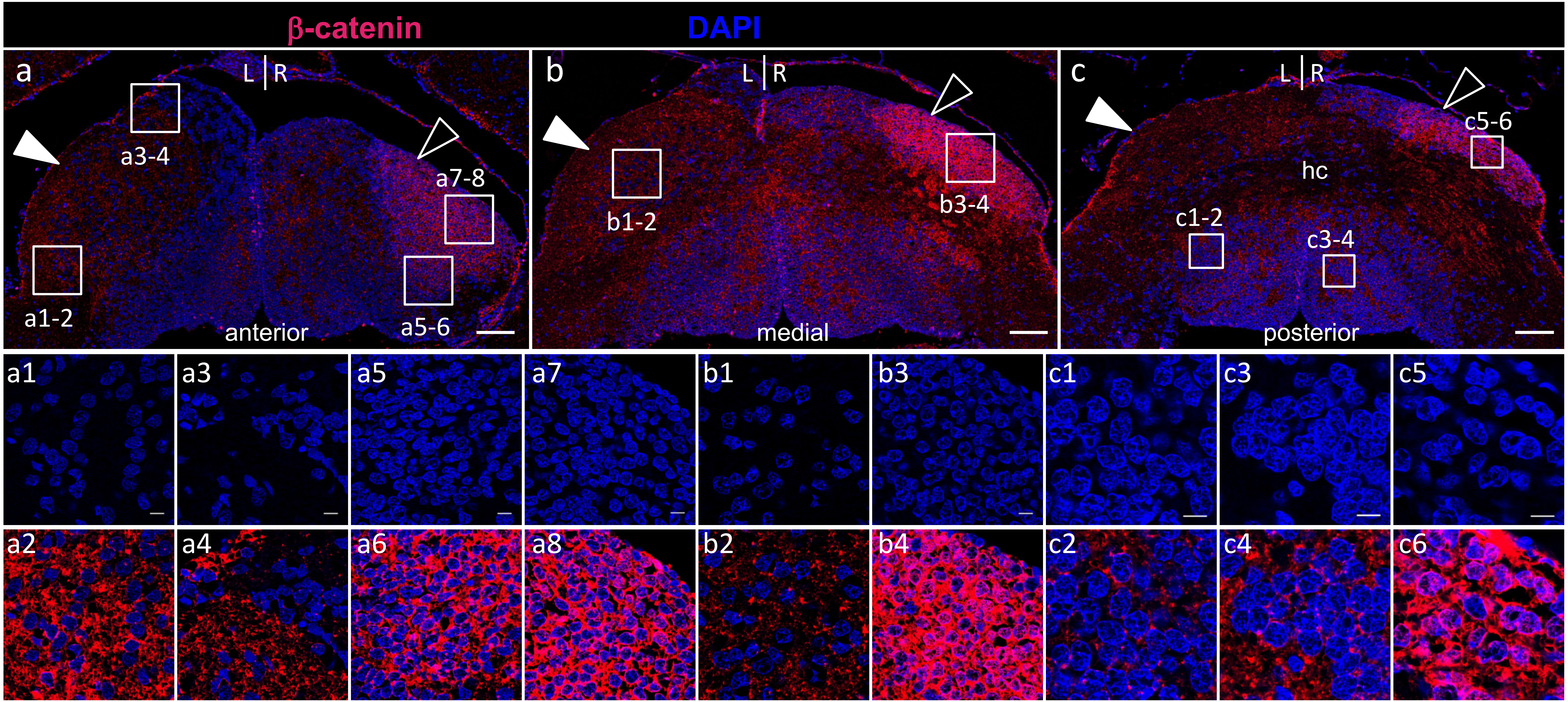


Supplementary Figure 17
